# Leveraging epigenomes and three-dimensional genome organization for interpreting regulatory variation

**DOI:** 10.1101/2021.08.29.458098

**Authors:** Brittany Baur, Jacob Schreiber, Junha Shin, Shilu Zhang, Yi Zhang, Mohith Manjunath, Jun S. Song, William Stafford Noble, Sushmita Roy

## Abstract

Understanding the impact of regulatory variants on complex phenotypes is a significant challenge because the genes and pathways that are targeted by such variants are typically unknown. Furthermore, a regulatory variant might influence a particular gene’s expression in a cell type or tissue-specific manner. Cell-type specific long-range regulatory interactions that occur between a distal regulatory sequence and a gene offers a powerful framework for understanding the impact of regulatory variants on complex phenotypes. However, high-resolution maps of such long-range interactions are available only for a handful of model cell lines. To address this challenge, we have developed L-HiC-Reg, a Random Forests based regression method to predict high-resolution contact counts in new cell lines, and a network-based framework to identify candidate cell line-specific gene networks targeted by a set of variants from a Genome-wide association study (GWAS). We applied our approach to predict interactions in 55 Roadmap Epigenome Consortium cell lines, which we used to interpret regulatory SNPs in the NHGRI GWAS catalogue. Using our approach, we performed an in-depth characterization of fifteen different phenotypes including Schizophrenia, Coronary Artery Disease (CAD) and Crohn’s disease. In CAD, we found differentially wired subnetworks consisting of known as well as novel gene targets of regulatory SNPs. Taken together, our compendium of interactions and associated network-based analysis pipeline offers a powerful resource to leverage long-range regulatory interactions to examine the context-specific impact of regulatory variation in complex phenotypes.

## Introduction

Genome-wide association studies (GWAS) have identified a large number of variants associated with different phenotypes and diseases. Approximately, 93% of all GWAS variants are located in non-coding regions and many are far away from any gene (Maurano et al., 2012). Understanding the mechanism by which such variants contribute to phenotypic variation is a significant challenge because the target gene(s) and downstream pathways of non-coding variants are unknown for many phenotypes and because of the cell type-specificity of such interactions between genes and variants. Furthermore, computational tools to identify how regulatory variants can impact molecular networks in a cell line-specific way are limited. Recent studies have shown that regulatory sequences such as enhancers can harbor non-coding variants that impact gene expression (Boyd et al., 2018; Corradin et al., 2014; Y. Zhang et al., 2018). Three-dimensional organization of the genome enables long-range regulatory interactions between enhancers and genes hundreds of kilobases apart through chromosomal looping that brings the enhancer in spatial proximity to target genes. Consequently, long-range interactions are emerging as an important component for the interpretation of regulatory variation (Cavalli et al., 2019; Pradhananga et al., 2020; Williams et al., 2020) and have been implicated in many diseases such as autoimmune diseases (Javierre et al., 2016) and cancer (Y. Zhang et al., 2019). Therefore, to systematically examine the impact of regulatory variants, we need long-range regulatory interaction maps across multiple cell lines as well as computational tools that can leverage these interactions to identify the gene networks that are targeted by sets of regulatory variants.

Genome-wide Chromosome Conformation Capture (3C) technologies, such as Hi-C (Kempfer & Pombo, 2020), offer comprehensive/genome-wide maps of 3D proximity of genomic loci from which we can identify long-range regulatory interactions. However, detecting long-range interactions between regulatory elements and genes requires high-resolution Hi-C datasets (e.g. 5kbp), which is limited to a handful of well-characterized model cell lines due to sequencing costs and the required number of cells for making reliable measurements at high resolution. A number of computational approaches have leveraged one-dimensional regulatory signals such as histone modifications, transcription factor (TF) binding and accessibility to predict long-range gene regulation using classification (Q. Cao et al., 2017; S. Roy et al., 2015; Whalen et al., 2016) and regression approaches (Belokopytova et al., 2020; S. Zhang et al., 2019). However, prediction of counts and interactions in new cell lines is still difficult and requires a large number of datasets for training that may not be available for many other cell lines and tissues. A related challenge is examining how regulatory variants impact different molecular networks. Computational approaches to identify molecular networks that are impacted by sequence variants have largely focused on coding variants (Eilbeck et al., 2017). However, incorporating regulatory variants requires us to link them to genes, which in turn requires comprehensive maps of long-range regulatory interactions across multiple cell lines.

To address both these challenges: the lack of comprehensive cell line-specific long-range regulatory interactions and defining the target pathways of a set of regulatory variants, we developed a computational pipeline comprising, L-HiC-Reg (Local HiC-Reg) to predict long-range interactions, and, graph-diffusion and multi-task graph clustering to identify the gene networks targeted by a set of variants (**Figure 1**). L-HiC-Reg is a Random Forests regression method that adapts our previous tool, HiC-Reg (S. Zhang et al., 2019) by leveraging the local structure of genomic segments to improve generalizability across cell lines and tissues. We used L-HiC-Reg models trained on high resolution Hi-C data to generate a compendium of predictions for 55 publicly available cell lines using real and imputed data from the Roadmap epigenomics database (Roadmap Epigenomics Consortium et al., 2015). Compared to existing resources such as GeneHancer (Fishilevich et al., 2017) and JEME (Q. Cao et al., 2017), our compendium is cell line-specific and is based on regression, rather than classification (JEME). The advantage of regression approaches is that the outputs can be used to identify larger organizational units such as topologically associating domains (TADs) as well as for detecting significant interactions, which can be both affected by a sequence variant. Our multi-task graph clustering approach identifies gene subnetworks for a particular phenotype of interest in a cell-line specific manner. Previous approaches to link non-coding SNPs to target genes have focused solely on the direct target genes using predicted or measured chromatin loops (Nasser et al., 2021). Our approach integrates both distal and proximal interactions to identify downstream pathways that may be disrupted as a result of a set of SNPs. Our predicted long-range interactions are corroborated with experimentally derived interactions from complementary ChIA-PET and Capture-Hi-C experiments and able to more comprehensively link sequence variants to genes compared to existing approaches.

**Figure 1.**
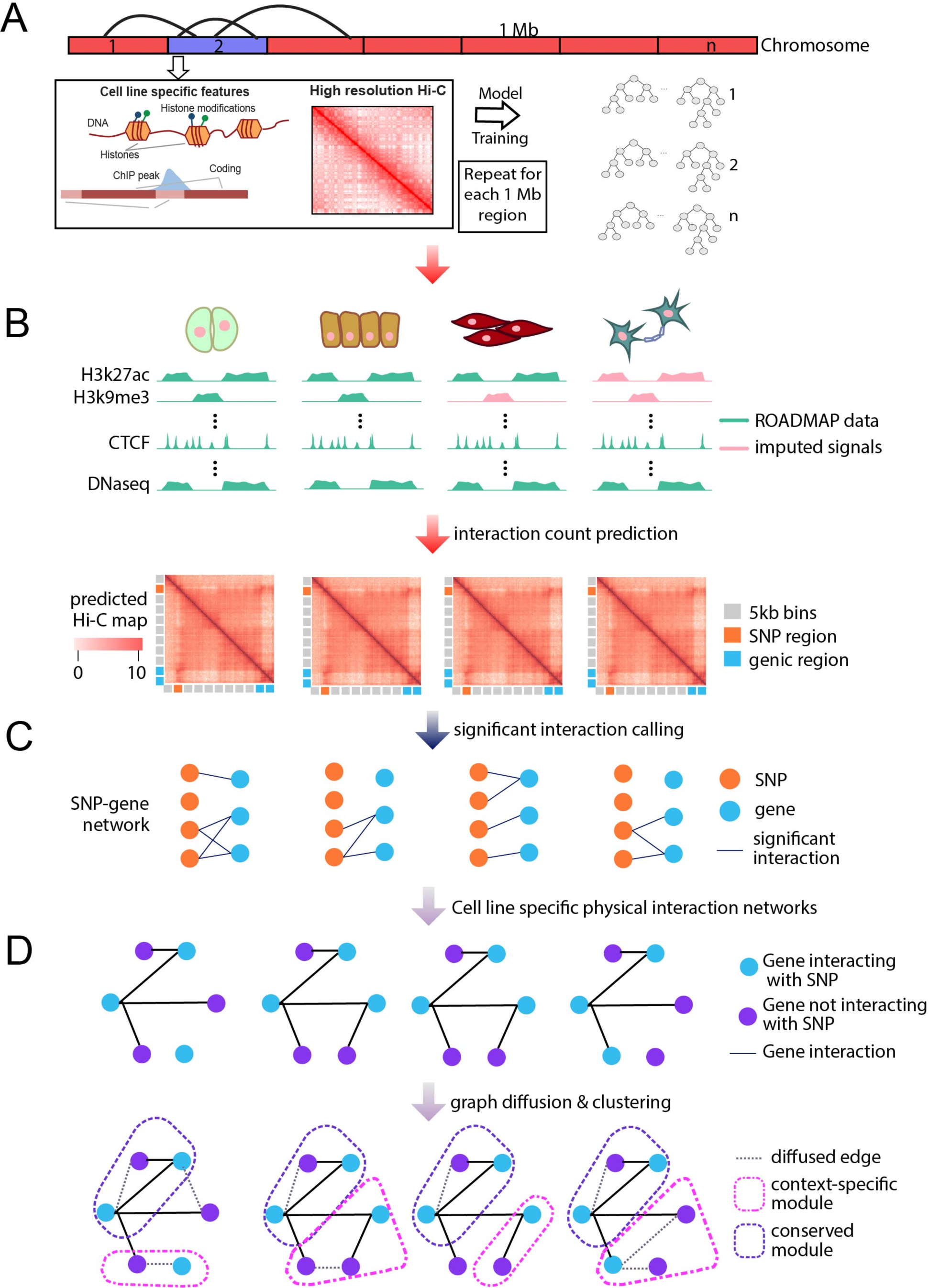
Overview of long-range interaction prediction and graph-based variant interpretation. **A.** L-HiC-Reg is trained on 1 MB regions of a chromosome with one-dimensional regulatory genomic datasets and high resolution Hi-C data using a random forest algorithm. **B.** The trained models are then applied to measured and imputed datasets in the Roadmap epigenomics database to generate a compendium of predictions in 55 cell lines. Generic 5kb bins are shown in gray, bins overlapping SNPs are in orange and TSS bins are in blue. **C.** SNPs and genes are connected in the SNP-gene network via long-range interactions with the SNPs for a given phenotype. Genes are scored based on the average significance of interactions with SNPs **D.** Physical interaction networks from protein-protein interactions and TF-gene interactions are used to perform graph diffusion on the scores from **C.** Multi-task graph clustering is used to identify gene subnetworks jointly across each cell line to identify pathways affected by the set of SNPs.

We applied our approach to 15 different phenotypes from the NHGRI-EBI GWAS catalog that represent different autoimmune, cardiovascular, neurological and cancer diseases (MacArthur et al., 2017). We identified gene subnetworks that are enriched with specific SNPs and exhibit network rewiring across the 55 cell lines. These subnetworks connect known and novel genes associated with a phenotype of interest offering novel hypotheses and prioritization of validation experiments needed to improve our understanding of a specific phenotype.

## Results

### L-HiC-Reg accurately predicts contact counts in a large number of cell lines

We previously developed HiC-Reg, an approach to computationally predict contact counts based on one-dimensional signals such as histone marks and transcription factor binding sites (S. Zhang et al., 2019). HiC-Reg performs well across chromosomes, however predicting across cell lines is still challenging and requires a large number of one-dimensional signals. We hypothesized that a single, global Random Forests regression model may not be able to capture locus-specific information for predicting interaction counts from one-dimensional signals. Therefore, we developed L-HiC-Reg, a “local” version of HiC-Reg that segments the chromosome and trains a separate model on each segment, as opposed to training on the whole chromosome (**Figure 1A**, **Methods**). L-HiC-Reg also uses a smaller number of one-dimensional signals, available for many cell lines, than originally used for HiC-Reg. Briefly, we used features from 7 experimentally measured and 3 computationally derived one-dimensional signals, together with genomic distance to represent a pair of regions. These datasets include the architectural protein CTCF, repressive marks (H3k27me3, H3k9me3), a mark associated with active gene bodies and elongation (H3k36me3), enhancer specific marks (H3k4me1, H3k27ac), an activating mark (H3k4me3), cohesion component (RAD21), a general transcription factor (TBP) and DNase 1 (open chromatin). The histone marks were selected because they were the most frequently measured across the cell lines in the Roadmap database, which we used to generate our compendium of predictions (**Supp. Figure 1**). Datasets for CTCF, TBP and RAD21 were computationally derived from sequence and open chromatin (**Methods**) (Sherwood et al., 2014). Similar to HiC-Reg, L-HiC-Reg predicts contact counts between 5kb pairs using the one-dimensional regulatory signals such as histone marks, accessibility and architectural protein binding motifs. A pair of regions in L-HiC-Reg and HiC-Reg is represented by the features for both 5kb regions, as well as the average signals between the two regions (window signal) as initially proposed in TargetFinder (Whalen et al., 2016). We trained models using high-resolution (5kb) Hi-C datasets from Rao et al (**Methods**, Rao et al., 2014). We applied L-HiC-Reg to trained models on adjacent 1Mb regions and predict in the same 1MB region in a different cell line. We then concatenated the predictions for the entire chromosome.

To test whether L-HiC-Reg recapitulates contact counts more accurately than HiC-Reg across cell lines, we trained L-HiC-Reg and HiC-Reg models on the five 5kb-resolution Hi-C data sets from Rao et al. We then generated all possible cross-cell line predictions for each chromosome resulting in 440 total predictions for each method. We assessed the performance of L-HiC-Reg and HiC-Reg using distance stratified Pearson’s correlation computed on the pairs from the test cell line. The distance stratified correlation measures the correlation between predicted and true counts for genomic pairs at a particular distance. Distance stratification is important due to the high dependence of contact count on genomic distance. We take the area under the distance stratified correlation curve (AUC) as a measure of cross-cell line prediction accuracy. We find that in 270 of the 440 cross-cell line predictions, the AUC is higher for L-HiC-Reg than HiC-Reg (Kolmogorov-Smirnov (KS) p-value=0.0011). This suggests that in the majority of cross-cell line predictions, L-HiC-Reg recapitulates contact counts better than HiC-Reg (**Figure 2A**). Additionally, we compared the performance of L-HiC-Reg to a baseline model in which the count from the training cell line is used as the predicted count for the test cell line (“Transfer count”). L- HiC-Reg performs better than the transfer count for 275 of the 440 cross-cell line predictions (KS p-value=1.5431e-04), suggesting that L-HiC-Reg is predicting more cell-type specific contact counts (**Figure 2A**).

**Figure 2.**
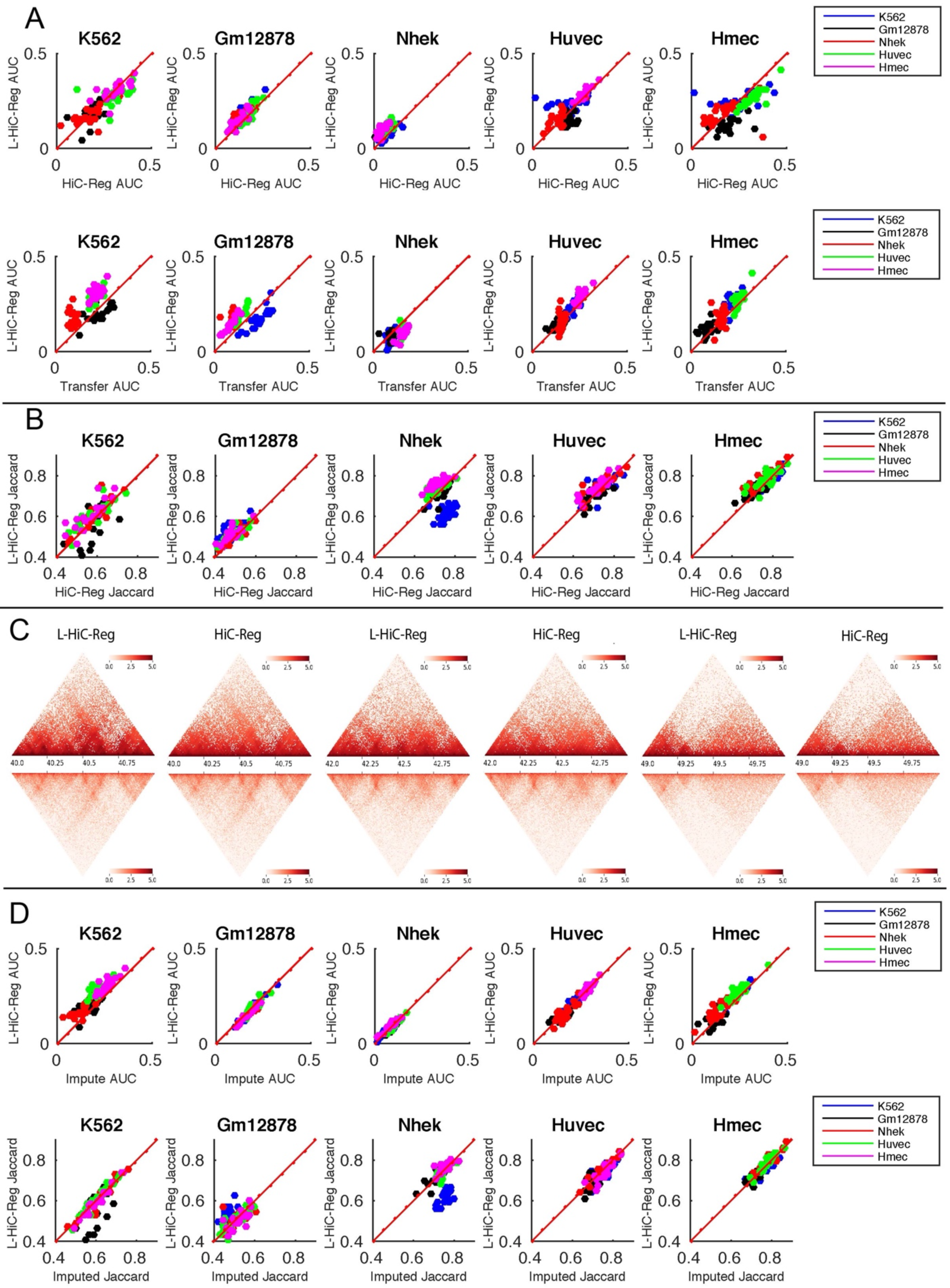
Evaluation of L-HiC-Reg predictions. **A.** Performance of L-HiC-Reg for count prediction between pairs of regions is assessed with the Area Under the correlation Curve (AUC) against HiC-Reg (top) and against transfer count (bottom). Each panel is a different test cell line and the color indicates the training cell line. **B.** Performance of L-HiC-Reg based on identified TADs from L-HiC-Reg predictions versus HiC-Reg predictions based on Jaccard coefficient similarity of TADs from true and predicted counts. Jaccard coefficient was used to assess the overlap of TADs found on the true counts with TADs found on the predicted counts by L-HiC-Reg (y-axis) and HiC-Reg (x-axis). **C.** Heatmaps of exemplar regions comparing Huvec L-HiC-Reg predictions and HiC-Reg predictions (top) against measured Huvec Hi-C data (bottom). **D.** Assessing predictions from measured versus imputed marks. AUC for L-HiC-Reg predictions generated from real one-dimensional datasets (y-axis) and imputed one-dimensional datasets (x- axis) in the test cell line (top). Jaccard coefficients comparing TAD recovery from measured (y- axis) and imputed data (x-axis) (bottom).

We next used the predicted counts from HiC-Reg and L-HiC-Reg to define topologically associating domains (TADs) by applying the Directionality Index (DI) TAD finding method (Sherwood et al., 2014). We compared these TADs to TAD identified by DI on the true counts based on a metric derived from the Jaccard coefficient (**Methods**). The Jaccard coefficient is a number between 0 and 1, with 0 representing no agreement and 1 representing perfect agreement. We computed the Jaccard coefficients for all cross-cell line predictions on all chromosomes for L-HiC-Reg and HiC-Reg (**Figure 2B**). We find that the Jaccard coefficients are higher in L-HiC-Reg than HiC-Reg in 302 of 440 cross-cell line predictions and is significantly better in two of the five cell lines (KS-test <0.05) and is comparable in the other cell lines. This suggests that L-HiC-Reg recapitulates TADs better than HiC-Reg when predicting in a new cell line. Visual inspection of the predicted matrices from the two methods for a 1MB region in the HUVEC cell line (**Figure 2C**) also demonstrated a clearer TAD structure when using L-HiC-Reg than HiC-Reg. Taken together these results show that L-HiC-Reg can accurately predict Hi-C contact count matrices in new cell lines.

### Prediction of Hi-C contact counts across multiple cell lines with publicly available epigenomic datasets

We next sought to apply L-HiC-Reg models on publicly available epigenomic datasets for different cell lines to create high-resolution *in silico* Hi-C maps for these cell lines. Here, we considered cell lines and tissues in the Roadmap Epigenome database for which a number of one-dimensional signals were available. These include DNase I for accessibility and six histone marks that are commonly measured (H3k27me3, H3k9me3, H3k36me3, H3k27ac, H3k4me1, H3k4me3), although often not within the same cell line. In particular, only 22 of the total 183 cell lines had all six epigenomic signals (**Supp Figure 1**). As the epigenomic dataset availability varies across cell lines, it can be difficult to make predictions from a model trained on features generated from a certain set of epigenomic marks that are not available in the test cell line. In order to consider as many cell lines as possible with L-HiC-Reg, we applied Avocado (Schreiber et al., 2020) a epigenomic signal imputation algorithm that combines tensor factorization and deep neural networks to produce an epigenomic signal prediction. The quality of the imputation improves as the number of datasets and cell lines increases. Therefore, we used the 5kb signals from 33 different datasets and 104 different cell lines from Roadmap, including the five cell lines from Rao et al. We used Avocado to generate epigenomic signals in 55 cell lines for which DNase I accessibility was available. In addition, we included the 5 Rao et al. cell lines to evaluate predictions of contact counts from imputed signals. We chose not to use imputed DNase I because we also use DNase I to generate the motif features for which a BAM file is needed and is needed downstream for identifying molecular networks affected by non-coding variants (**Methods**).

We applied our trained L-HiC-Reg models on measured and imputed signals, DNase I and additional TF features derived from accessibility to all 55 cell lines and the 5 Rao et al cell lines where all the marks were imputed. We wanted to determine if we could accurately predict contact counts in a test cell line where all histone features are derived from imputed datasets (worst case scenario). We generated all cross-cell line predictions for the 5 Rao et al. cell lines on all chromosomes using the imputed features in the test cell line, using the models trained on the original measured values on the train cell line. Based on the area under the distance stratified correlation, the resulting predictions were not significantly different from those generated from entirely real features (**Figure 2D**; KS test p-value=0.11). Additionally, we used the predictions based on imputed features and called TADs. The Jaccard coefficients from these TADs were also similar to the ones generated from real features (**Figure 2D,** KS test p-value=0.84). Taken together, these results suggest that there is not a significant deterioration in the quality of the predictions if imputed features are used.

### L-HiC-Reg long-range interactions are validated by complementary experimental sources and associated with highly expressed genes

We identified long-range interactions defined by significantly interacting regions (loops) using a distance stratified Binomial test (Duan et al., 2010) on the L-HiC-Reg predicted contact count matrices for each of the 55 cell lines. Across 55 cell lines, we identified between 681,889 and 1,481,911 significant interactions with an average of 983,250 interactions (q-value <0.05). To evaluate these significant interactions, we compared them against interactions identified from complementary experimental assays, ChIA-PET and Capture-Hi-C (**Methods**). We obtained 10 published ChIA-PET datasets for different factors (RNA PolII, CTCF and RAD21) and histone marks in multiple cell lines (Heidari et al., 2014; G. Li et al., 2012). We estimated fold enrichment of the ChIA-PET interactions with the significant interactions in each of the 55 cell lines (**Figure 3A**). We did the same for two other computationally-predicted resources, JEME (Q. Cao et al., 2017) and GeneHancer (Fishilevich et al., 2017). GeneHancer is a network of enhancer-promoter interactions that scores potential enhancers and their interactions based on evidence from various publicly available datasets, such as eQTL, functional annotation and chromatin accessibility. JEME is a classification approach that incorporates information from multiple cell lines to predict an interaction between a promoter and candidate enhancers. ChIA-PET interactions were all enriched (fold enrichment > 1) in the L-HiC-Reg predictions (**Figure 3A**). Additionally, L-HiC-Reg had a higher fold enrichment compared to JEME and GeneHancer in 7 out of 10 datasets.

**Figure 3.**
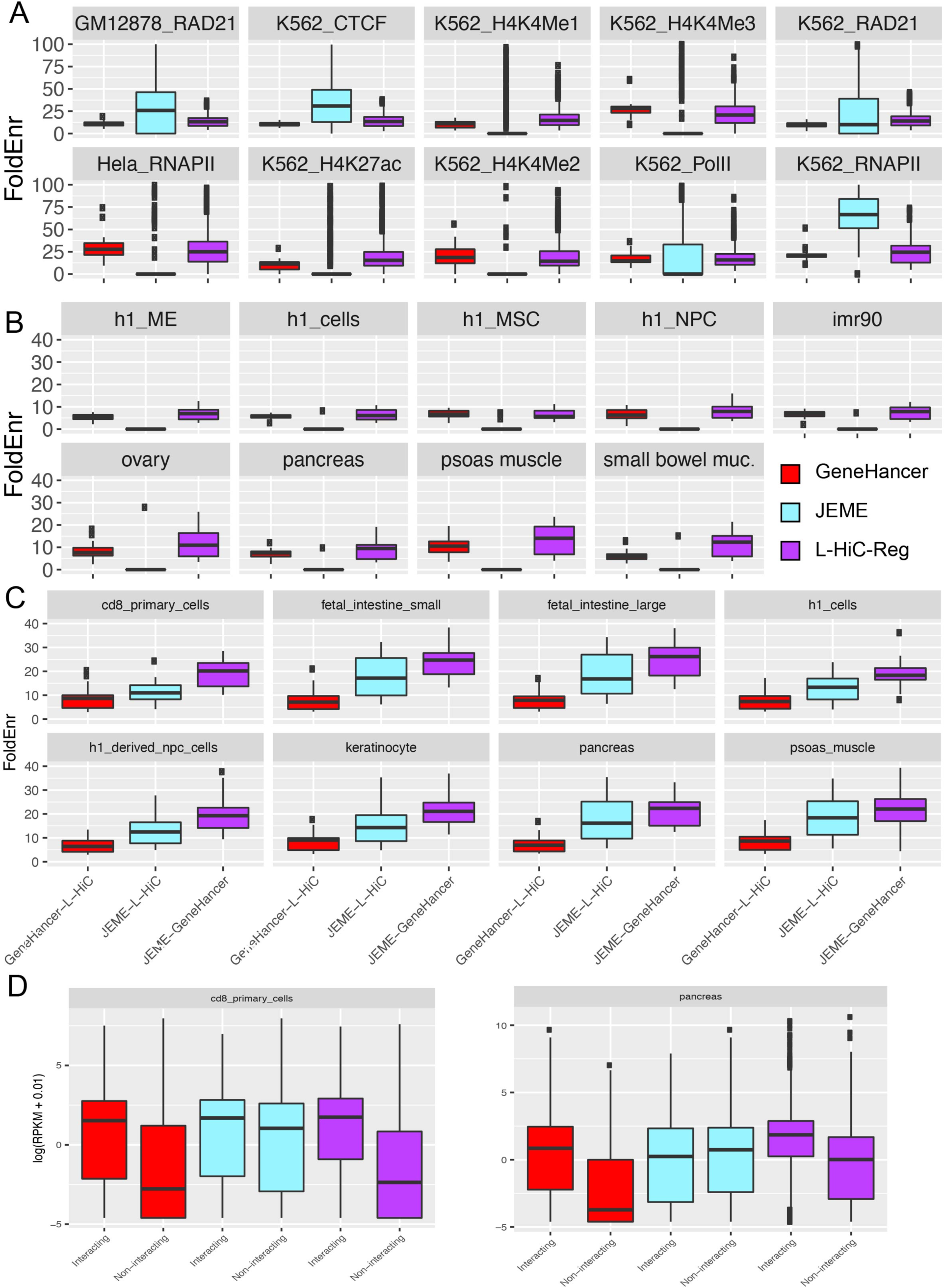
Performance of long-range interaction prediction approaches against experimentally derived datasets. **A.** Fold enrichment of JEME (cyan), GeneHancer (red) and L-HiC-Reg (purple) against gold standard ChIA-PET datasets. Predictions for each cell line from L-HiC-Reg and JEME were compared against each ChIA-PET dataset shown as a separate panel. GeneHancer is not cell line specific and had single set of predictions that were compared against all ChIA-PET datasets. Enrichment was calculated separately for each chromosome. The box plot shows the distribution of the enrichment values for each chromosome for a pair of predicted and ChIA-PET interactions **B.** Fold enrichment against capture Hi-C data. The predictions from L-HiC-Reg and JEME were matched to the capture Hi-C dataset based on cell line. Enrichment was calculated separately for each chromosome. **C.** Comparing the fold enrichment of pairs of computational approaches GeneHancer and L-HiC-Reg (red), JEME and L-HiC-Reg (cyan) and JEME and GeneHancer (purple) in matched cell lines. **D.** Expression of genes (RPKM) with interactions compared to genes with no interaction in GeneHancer (red), JEME (cyan) and L-HiC-Reg (purple). L-HiC-Reg and JEME were matched for the cell line.

We next compared L-HiC-Reg, JEME and GeneHancer predictions to several Capture Hi-C datasets from Jung et al., 2019, which profiled 27 cell lines, 9 of which overlapped with the cell lines we examined (Jung et al., 2019). We compared directly with the matching cell type for all chromosomes. For GeneHancer, which does not predict cell line specific interactions we considered the same set of interactions for each cell line. L-HiC-Reg interactions exhibit the highest fold enrichment for these datasets compared to JEME and GeneHancer (**Figure 3B**).

We also compared the predicted interactions from each of the three methods, L-HiC-Reg, Genehancer and JEME, to each other. All computational methods were mutually enriched among each other with L-HiC-Reg exhibiting the highest enrichment in JEME interactions (**Figure 3C**). This is likely because L-HiC-Reg and JEME are both cell-line specific and random forest-based. The fact that there is a high enrichment among the different computational approaches is encouraging, especially since GeneHancer has a different underlying model.

Chromosomal looping is often associated with changes in gene expression by bringing enhancers bound by transcription factors in close proximity to the promoters of their target genes. TRIP (Thousands of Reporters Integrated in Parallel) is an experimental technique that inserts barcoded transgene reporters randomly into the genome and can measure transcriptional activity (Akhtar et al., 2013). It has previously been demonstrated that transgenes mapped to promoter-interacting fragments demonstrate higher expression values over all genomic distances (Cairns et al., 2016). As another validation, we linked significant interactions to genes in either or both of the interacting regions. Across the four cell lines that were common to JEME and L-HiC-Reg and had RNA-seq data available, L-HiC-Reg linked an average of 10,341 genes over all the cell types, and such genes had a significantly higher expression (t-test P-value < 0.05) than genes that are not associated with significant interactions (**Figure 3D**, **Supp Figure 2**). GeneHancer linked 14,467 genes and also had a large difference between the genes with interactions versus genes that did not have interactions (t-test P-value<0.01). JEME linked an average of 5,810 genes across the cell types and the difference in expression between genes with and without interactions was significant, likely because of the small number of genes linked by JEME. Taken together these results show that our compendium of L-HiC-reg interactions are well supported by existing complementary assays and are of as good or higher quality than other computational predictions.

### Leveraging L-HiC-Reg long range interactions to link regulatory SNPs to genes across diverse cell lines

We next used our compendium of significant L-HiC-Reg interactions across 55 cell lines and tissues and tissues to interpret regulatory variation by linking non-coding SNPs to genes. We downloaded the NHGRI-EBI GWAS catalog, which contains curated GWAS SNPs for thousands of phenotypes (MacArthur et al., 2017). For a given GWAS, we defined a non-coding SNP as one that was intergenic and identified all SNPs in Linkage Disequilibrium (LD) with it, resulting in a total of 23,116 GWAS non-coding SNPs and 364,893 SNPs in LD (**Methods**). We mapped all non-coding SNPs and SNPs in LD to genes using L-HiC-Reg significant interactions (**Methods**). Of the total 23,116 GWAS SNPs we were able to map 35.16% to genes across our 55 cell lines with an average 7,450 genes mapped in any cell line with ∼30k interactions **(Figure 4A**). In comparison, both JEME and GeneHancer mapped fewer proportions of SNPs (24.47% with JEME and 28.4% using GeneHancer). Similarly, for the SNPs in LD, L-HiC-Reg mapped a higher proportion to genes (27.19%, **Supp. Figure 3**) compared to JEME (18.7%) or GeneHancer (21.17%).

**Figure 4.**
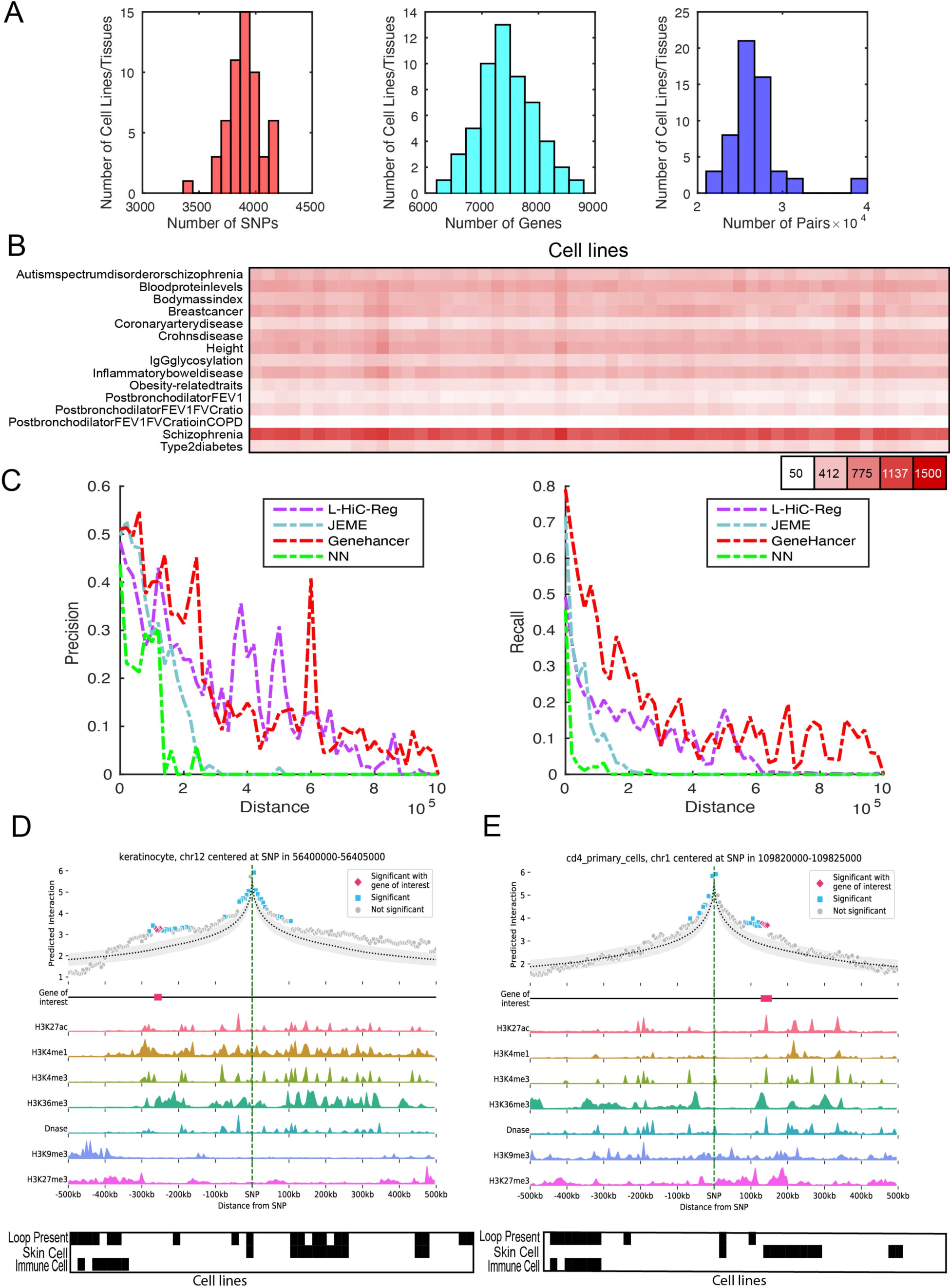
**A.** Distribution plots for all GWAS non-coding SNPs and SNPs in LD that were linked to genes across the 55 cell lines (left), genes linked to SNPs (middle) and SNP-Gene pairs (right). **B.** Number of genes associated with SNPs across phenotypes (rows) and cell lines (columns). **C.** Precision (left) and recall (right) of L-HiC-Reg, JEME, GeneHancer and nearest neighbor (NN) for eQTL SNP-Gene associations. **D.** Contact count and one-dimensional signals centered around the vitiligo-associated SNP rs10876864. The *GDF11* gene is highlighted on the Gene of interest track. **E.** Contact count and one-dimensional signals for coronary artery disease-associated SNP rs599839. The *PSMA5* gene is highlighted on the Gene of interest track. Presence-absence pattern of the SNP-gene interaction is shown using the black and white matrix with columns corresponding to cell lines.

As SNP-gene associations could indicate regulatory relationships between the SNP and the gene by impacting its expression, we leveraged expression QTL (eQTL) datasets from the GTEX database (Lonsdale et al., 2013) to assess the SNP-gene associations from each of the methods (**Methods**). We considered 4149 GWAS and 73662 LD SNPs for 15 phenotypes in the GWAS catalog with the largest number of SNPs (**Table 1**). The number of genes linked by L-HiC-Reg varied between 100-2400 genes across cell lines for any given phenotype. The exception was Schizophrenia for which we were able to map many more genes likely because of more SNPs (**Figure 4B**). For each set of SNP-gene associations, we calculated the proportion of predicted interactions that are also supported by eQTL SNP-gene associations (precision) and the proportion of eQTL SNP-gene associations in each predicted set (recall). As a baseline, to assess the importance of leveraging long-range interactions to link SNPs to genes, we also found the closest gene (nearest neighbor, NN) based on genomic distance to each GWAS and LD SNP and compared these SNP-gene associations to eQTL associations. To account for the fact that eQTL SNP-gene pairs are more likely to be close together and many of the predictions are between close regions, we stratified these two quantities by distance **(Figure 4C**). SNP-associations from L-HiC-Reg and GeneHancer have higher precision than JEME for all distance bins compared, and GeneHancer outperforms L-HiC-Reg for more distance bins (35 vs 16). Based on recall, GeneHancer was generally better than L-HiC-Reg with the exception of associations at 400-600k. Both methods were better than JEME for recall. The higher performance of GeneHancer is not surprising as it uses GTEx data, along with many other resources, to assign a score to an interaction. JEME performs well in close-range interactions in precision and recall, but rapidly declines in precision and recall for longer-range interactions. Notably, all computational approaches perform better than the NN approach and, for distances greater than 200kb, very few NN genes are in eQTL (**Figure 4C**). Taken together, these results highlight the importance of leveraging long-range interactions to link SNPs to genes, particularly those predicted at >200kb distances.

**Table 1.** Statistics for 15 phenotypes analyzed using our framework. Shown are the total number of GWAS non-coding SNPs, total number of SNPs in LD with these SNPs, total number of SNPs (GWAS and LD) linked to genes, total number of genes linked to SNPs and total number of SNP- gene pairs.

| Phenotype | GWAS | LD | SNPs<br>Mapped | Genes<br>mapped | Pairs |
| --- | --- | --- | --- | --- | --- |
| Autism spectrum disorder or schizophrenia | 134 | 300 | 365 | 808 | 15439 |
| Blood protein levels | 154 | 2837 | 1975 | 1192 | 37676 |
| Body mass index | 467 | 10791 | 2533 | 1294 | 47762 |
| Breast cancer | 354 | 1697 | 644 | 1641 | 8074 |
| Coronary artery disease | 235 | 174 | 193 | 1036 | 1737 |
| Crohn's disease | 170 | 6171 | 3181 | 1232 | 54566 |
| Height | 266 | 9530 | 3233 | 1451 | 37839 |
| IgG glycosylation | 223 | 4093 | 1266 | 760 | 28467 |
| Inflammatory bowel disease | 177 | 5113 | 2458 | 1349 | 34933 |
| Obesity-related traits | 319 | 2676 | 942 | 988 | 7094 |
| Postbronchodilator FEV1 | 301 | 3721 | 556 | 568 | 6868 |
| Postbronchodilator FEV1/FVC ratio | 579 | 10003 | 563 | 979 | 4219 |
| Postbronchodilator FEV1/FVC ratio in COPD | 137 | 3883 | 235 | 167 | 1216 |
| Schizophrenia | 397 | 9714 | 3331 | 2378 | 72364 |
| Type 2 diabetes | 236 | 2959 | 1040 | 813 | 11508 |

Several of L-Hi-C-Reg SNP-gene associations are supported by experimental assays targeting SNP-gene associations with Capture-Hi-C (Cavalli et al., 2019; Pradhananga et al., 2020). We obtained two large-scale Capture-Hi-C datasets that investigated GWAS SNP-gene interactions. One of these focused on blood cell traits and generated a Capture-Hi-C dataset in lymphoblastoid cells (Cavalli et al., 2019). The second study used Capture-Hi-C in macrophages and studied immune and cardiovascular risk variants (Pradhananga et al., 2020). In addition to Capture-Hi-C datasets, these studies used a number of computational filters, such as motif disruption, chromatin signal to identify SNP-gene associations. We filtered these SNP-gene associations based on whether interactions were distal which is defined as regions that interacted with designed promoters or other distal regions (Cavalli et al., 2019). One of these interactions from Cavalli et al., linked the SNP rs10876864 to the gene *GDF11*, based on Capture-Hi-C, motif disruption and chromatin signals. This SNP is associated with vitiligo, a skin disease, and *GDF11* is known to be involved in the regulation of skin biology. In our predictions, we found this SNP to loop to *GDF11* in several skin and immune cell lines (**Figure 4D**). The predicted count profile shows a significantly high count value for this SNP and GDF11, compared to the background count distribution in one exemplar cell line (keratinocyte, gray shading, **Figure 4D**). Furthermore, inspection of the one-dimensional signals in the keratinocyte cell line showed a clear association of H3k4me1, an active enhancer mark, between the SNP and the gene. Similarly, we found an interaction between the SNP rs599839, associated with cardiovascular diseases, and the gene *PSMA5*, that was present among immune cell lines (**Figure 4E**). Inspection of the predicted count profile and one-dimensional signals around the rs599839 SNP in cd4 primary cells again highlighted the high interaction count for this interaction supported by one-dimensional signals such as DNase I and H3k4me3. Overall, these results offer experimental support for our predictions many of which are likely valid regulatory interactions based on the overlap with eQTL studies.

### Identifying downstream pathways impacted by regulatory variants

Our results so far showed that we are able to successfully link regulatory variants to genes across diverse cell lines. However, phenotypic variation is typically driven by groups of genes that interact in a larger unknown pathway and therefore interpretation of regulatory variation requires us to identify the impact of sequence variants at the pathway level. Systematic identification of pathways affected by a set of SNPs in a specific cell line is challenging because not all pathways and their components are known and furthermore to what extent they are preserved across different cell and tissue types. To examine the impact of a set of variants at the pathway level, we developed a graph-theoretic framework based on graph diffusion and multi-task graph clustering to simultaneously define gene subnetworks representative of gene pathways across different cell lines (**Figure 5**). We first created a collection of cell line specific molecular interaction networks that combined protein-protein interaction networks, promoter proximal and distal transcription factor (TF)-gene interactions across each of the 55 cell lines (**Methods**). The protein-protein interactions were context-unspecific while the TF-gene interactions were cell line-specific. For a given phenotype, we scored a gene predicted to interact with a SNP based on the average - log(Pvalue) of its L-HiC-Reg interactions with SNPs (**Methods**). We used graph diffusion to define a set of direct and indirect gene hits (defined by the top 1% genes with no SNP interactions) of SNPs and created cell-line specific weighted graphs where the edge weight corresponded to the strength of a diffusion signal from one of the direct hits. We then applied a multi-task graph clustering approach, Muscari, to the graphs to identify different gene networks based on their connectivity to genes with SNPs (Shin et al., 2021). Our multi-task graph clustering approach takes as input a relationship tree between the cell lines and leverages these relationships while defining gene subnetworks for each cell line. We used the similarity of the H3K4me3 signal in gene promoters between pairs of cell lines and performed a hierarchical clustering to obtain this tree (**Supp Figure 3A**). The H3K4me3-based tree was similar to a tree based on shared long-range interactions between pairs of cell lines (**Supp Figure 3B**, **C)**. We applied our pipeline to 15 phenotypes in the GWAS catalog with the most non-coding SNPs (**Figure 4**, **Table 1**). The phenotypes include a range of neuronal (autism spectrum disorder and schizophrenia), cardio- vascular (coronary artery disease), and auto-immune disorders (Crohn’s disease) as well as other diseases (breast cancer, Type 2 diabetes). Below we discuss one of these phenotypes, Coronary Artery Disease. The results from our other phenotypes are available at https://github.com/Roy-lab/Roadmap_RegulatoryVariation.

**Figure 5.**
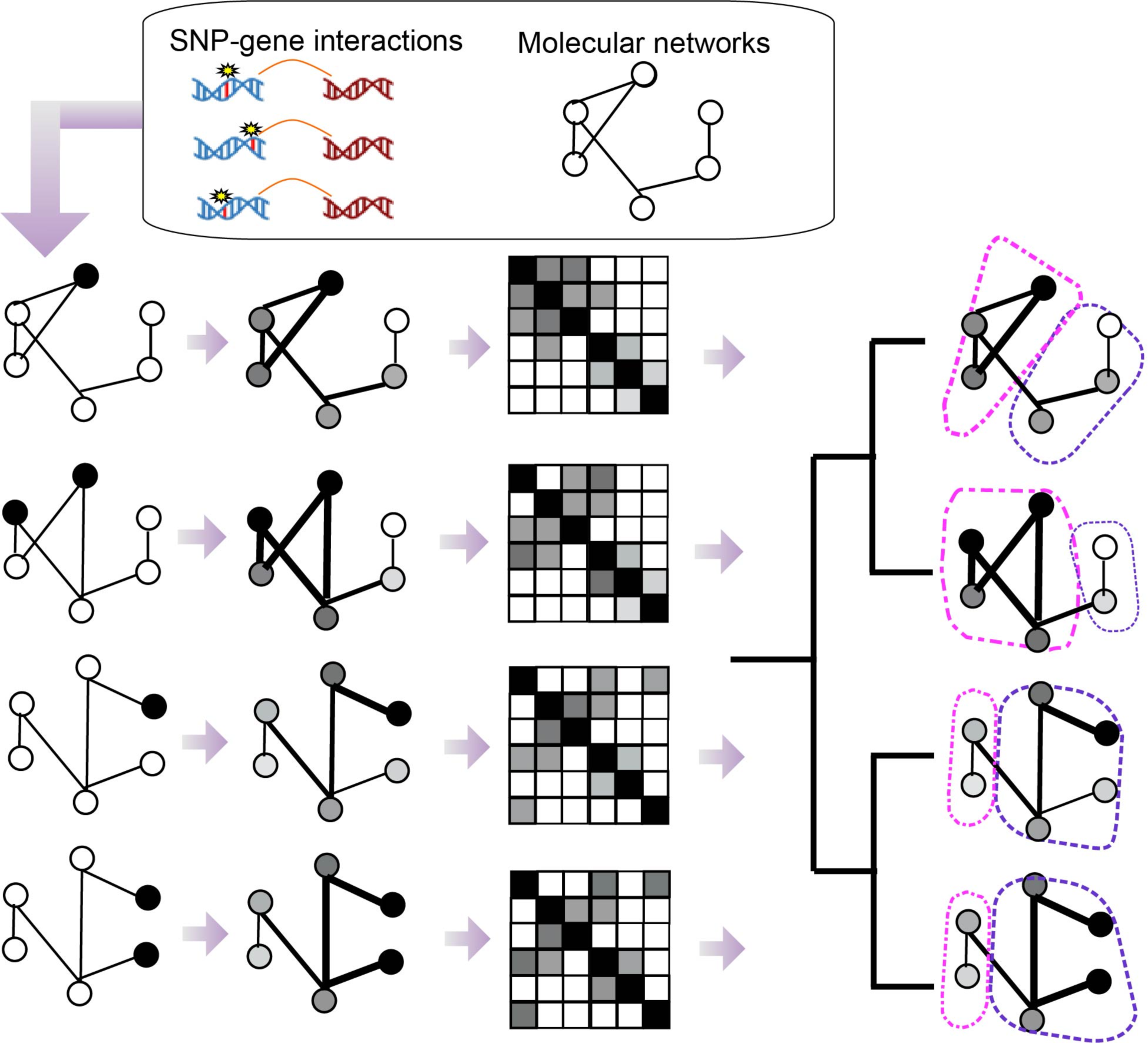
Overview of our Multi-task Graph Clustering (MTGC) approach. Genes are scored based on their significant interactions with SNP-containing regions and mapped to a physical interaction network. Graph diffusion is performed to obtain a fully connected adjacency matrix. The input into the the multi-task graph clustering approach is a matrix for each cell line, the number of clusters and relationship tree for the cell lines.

Coronary Artery Disease (CAD) is a type of cardio-vascular disease associated with thickening of arterial walls and is a major cause of disease in developed countries (Malakar et al., 2019). Recent GWAS of CAD has shown a significant portion of associated SNPs to lie in non-coding regions of the DNA (Selvarajan et al., 2021; Villar et al., 2020). Using L-Hi-C-Reg we mapped a total of 193 SNPs (141 GWAS and 52 in LD) of total 409 SNPs to 1036 genes across the 55 cell lines with a total of 1737 interactions (**Figure 6A, Table 1**). Among the cell lines that had the largest number of SNPs were fetal heart, skin and immune cell lines like CD4 and CD34 (**Figure 6B**). In contrast, muscle cell lines ranked lowest based on the number of genes. After graph diffusion and selecting the top 1% genes in each cell line, we had a total of 1866 genes across the cell lines (1449-1821 genes in any cell line). We tested the optimal number of clusters for Muscari based on modularity of clusters defined across the 55 cell lines and found k=7 to be best. Using Muscari, we were able to obtain cluster assignments for 1625 of the 1866 genes. We examined the clusters for enrichment of Gene Ontology (GO) terms in each of the 55 cell lines (**Supp Figure 5**, **Methods**). The clusters exhibited enrichment in diverse processes including cell cycle, transcription, and signaling **(Figure 6B, Supp Figure 5**). In particular, cluster 2 is enriched for terms relevant to CAD such as blood coagulation, wound healing, platelet activation and regulation of fluid levels (**Supp Figure 5**). Dysfunctional coagulation has been shown to be associated with coronary artery disease (H. Roy et al., 2009). Low mean platelet volumes are associated with worse outcomes for CAD patients (Wada et al., 2018). Additionally, heart is a top cell line for cluster 2 (**Supp Figure 5**).

**Figure 6.**
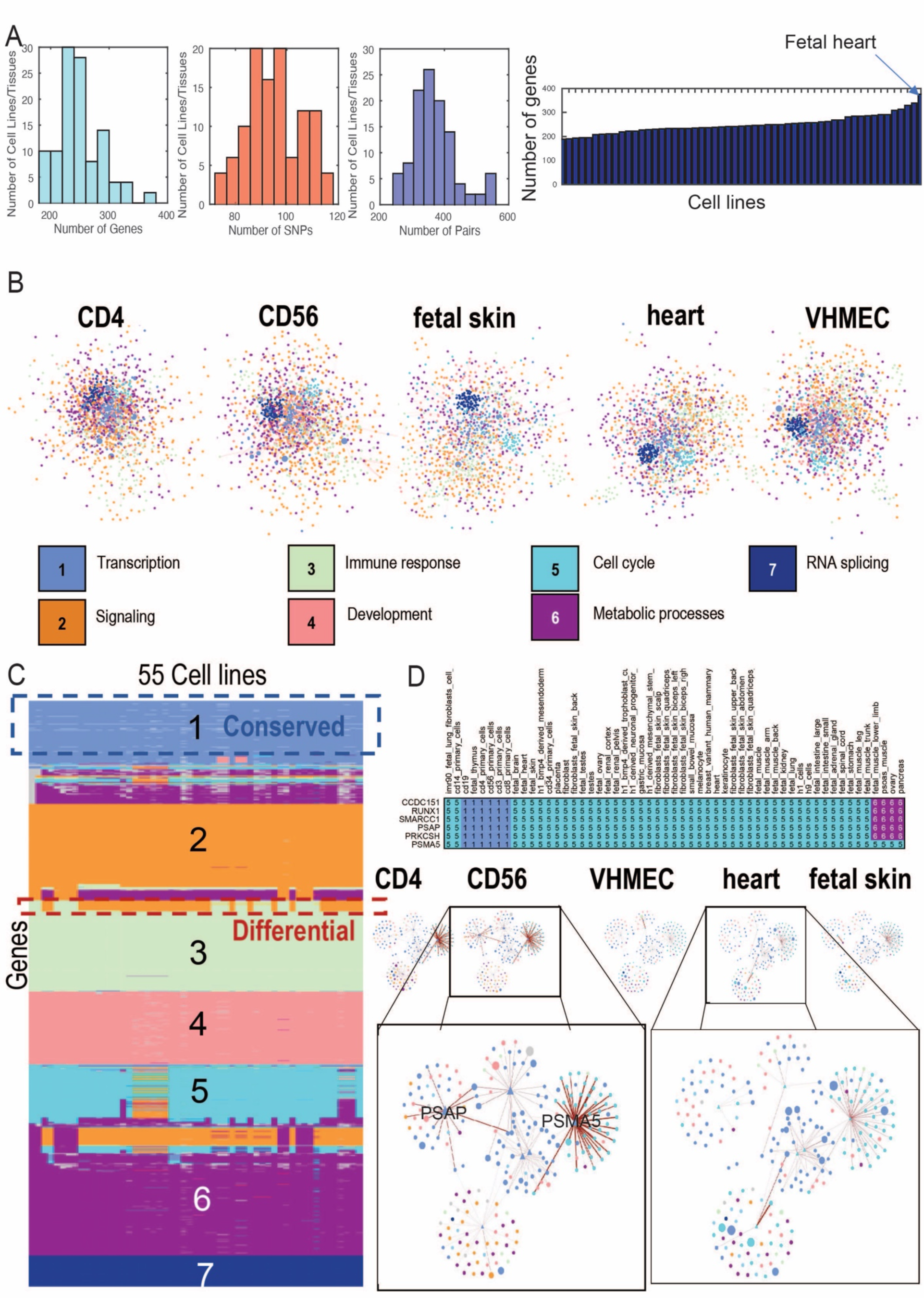
Interpreting regulatory variants in Coronary Artery Disease (CAD). **A.** Distribution of the number of genes linked to SNPs (cyan), SNPs linked to genes (red) and number of pairs (blue) across tissues (left). Number of genes linked to SNPs for every cell line (right). **B.** Example networks with nodes (genes) colored by their clustering assignment and the major GO enrichment process for each cluster. **C.** Cluster assignments across the genes (rows) and cell lines (columns). **D.** An example transitioning gene set. Rows are the genes and columns are the cell lines with the color indicating cluster assignment (top). Networks of the genes in this set (triangles) and their nearest neighbor target genes (circles) with edge weight represented by edge thickness (bottom).

We next examined the gene cluster assignment across the 55 cell lines to identify genes that change their cluster assignment across cell lines. Such genes are differentially connected across cell lines and could represent cell-line specific network components that are targeted by a set of SNPs. We clustered genes into gene sets based on their gene assignment across cell lines (**Methods**). Majority of the genes (1201 genes, 74%) did not change their assignment (**Figure 6C**). The remaining 424 genes that changed their cluster assignment and therefore differentially connected, fell into 17 gene sets. Most (12) of our gene sets were associated SNPs in at least one cell line indicating they were jointly targeted by these SNPs. We examined each of these gene sets based on the presence absence patterns SNP-gene interactions across each cell line and whether this corresponded with the change in cluster assignment. Three of the 15 gene sets, #37, #46, #53 showed concordant changes in cluster assignment and SNP-gene interaction (**Supp Figure 6**). Genes in set #46 were in cluster 1 in the immune cell types but transitioned to cluster 5 in most of the other cell lines (**Figure 6D**). This gene set included *PSMA5* and *PSAP* that had a number of CAD SNPs associated with them. This gene set exhibits differences in the diffused edge weights between immune cell lines and non-immune cell lines, particularly around the *PSMA5* gene (**Figure 6D**). The association of *PSMA5* with one of the SNPs was also identified by a Capture-HiC experiment (**Figure 4E**). *PSAP* is a membrane protein that plays a key role in lysosome function, and dysfunction of lysosomes (Zhou et al., 2015) has been associated with a number of cardiovascular diseases including CAD (Chi et al., 2020). *SMARCC1* is a chromatin remodeler, which is part of the SWI/SNF complex involved in the development of cardiovascular tissues (Vieira et al., 2017). While *PSMA5* has been shown before to associate with CAD GWAS SNPs, both *PSAP* and *SMARCC1* are novel predictions from our study. Gene set #37 is enriched in cell-matrix adhesion and integrin signaling. Cell adhesion molecules, including integrin, are biomarkers of CAD (Jang et al., 1994). This gene set additionally contains *COL7A1*, a collagen protein, which are abundant in the extracellular matrix and have been implicated in heart disease including CAD (Kunicki & Ruggeri, 2001). Gene set #53 contains a heat shock protein, *HSP90AB1* and *NUSAP1*, a gene involved in microtubule organization. Heat shock proteins have also been associated with heart failure (Ranek et al., 2018). Cytoskeleton alterations, including microtubules, have also been associated with heart disease (Hein et al., 2000; Scarborough et al., 2021). Taken together, our analysis of CAD illustrates the utility of applying our network-based interpretation framework to pinpoint specific gene pathways that could be targets of different regulatory SNPs. We identified both known and novel genes relevant to CAD and based on their biological roles represent plausible candidates for future validation studies.

## Discussion

Interpretation of non-coding variation and how it impacts phenotypic variation is a significant challenge in human genomics because of our limited knowledge of which genes and pathways these variants act upon. Long-range regulatory interactions between distal sequence elements and genes are a powerful mechanism by which variants can impact gene expression. However, high-resolution maps of such interactions are missing for most cell lines and biological contexts due to the cost of generating such Hi-C datasets. Furthermore, tools to interpret a set of variants to identify pathways that are impacted in a cell line specific manner are limited. Here we presented a computational pipeline to first generate *in silico* Hi-C maps across a large compendia of cell lines and then leverage these maps within a graph-theoretic framework to define gene subnetworks targeted by sets of sequence variants. Our pipeline comprises a L-HiC-Reg, a tool to use real and imputed signals from the Roadmap epigenomics database to predict high-resolution Hi-C maps across multiple cell lines and multi-task graph clustering to simultaneously define gene subnetworks across multiple cell lines. Using our approach, we were able to link a higher proportion of variants to genes compared to existing approaches, many of which are supported by auxiliary experimental sources such as eQTL, Capture-Hi-C and ChIA-PET datasets. We used our approach to predict cell line-specific pathways across 55 cell lines for 15 different phenotypes. With our graph-theoretic framework, we were able to find gene sets that change their connectivity across cell lines and harbor sequence variants for a phenotype.

Our computational pipeline and associated resources greatly expands on previous efforts for leveraging long-range interactions to link sequence variants to genes. In particular, compared to some resources (Fishilevich et al., 2017), our resource is context-specific comprising predicted loops and count matrices for individual cell lines . Second, our regression-based model, L-HiC-Reg predicts contact counts for every genomic pair, as opposed to methods which directly generate the set of interactions (Q. Cao et al., 2017). By predicting counts, we offer the end-user the flexibility to apply different statistical tools to define interactions and also identify higher order units of organization such as topologically associating domains (TADs). Finally existing approaches for interpreting regulatory variation using long-range interactions (Cavalli et al., 2019) typically operate at the level of individual SNP-gene pairs. Our pipeline leveraged cell line-specific regulatory interactions from our long-range interactions, accessibility, and cell line non-specific protein-protein interactions to predict gene subnetworks that targeted by sets of variants in a cell line-specific manner.

Functional validation of regulatory variant interactions to genes is challenging because of the large number of possible associations using both experimental and computational methods. By overlaying variants on a molecular network, with nodes and edges reweighted by their propensity to participate in long-range interactions with SNPs, we were able to narrow possible gene candidates that could be targeted by a SNP. We leveraged cell line-specificity of our networks to identify genes that change their clustering structure and therefore network connectivity across cell lines and found many of these gene sets to have sequence variants. Furthermore, correlating the change in network connectivity with the presence-absence pattern of the sequence variant-gene interaction helped narrow down the list of candidate gene-variant interactions that could be experimentally validated.

We used our new resource to examine non-coding variants in 15 different phenotypes spanning different disease and normal phenotypic traits. For each of the phenotypes we defined cell line specific gene targets and subnetwork across 55 different cell lines. Our predictions are available for download at https://github.com/Roy-lab/Roadmap_RegulatoryVariation. Comparison of the subnetworks across cell lines showed that most genes do not change their network connectivity. However, changes in network connectivity was often associated with their tendency to link to SNPs. One such example for the Coronary Artery Disease (CAD) that identified a set of 6 genes, that were predict to interact with CAD SNPs in immune cell types. We found a large number of connections with the genes *PSMA5*, *PSAP* and to a lesser extent, *SMARCC1* in the immune cell types, which were depleted in the other cell lines. While *PSMA5* was shown previously to link to CAD SNPs by a previous Capture-Hi-C study (Cavalli et al., 2019), *PSAP* and *SMARCC1* are novel to our study. Taken together, these results suggest that our approach can identify relevant gene subnetworks that are associated with SNPs for a phenotype of interest and can be used to investigate the effects of non-coding variation.

Our work can be extended in a number of ways. Here, we relied on regulatory interactions based on the presence of sequence-specific motifs, which did not include transcription factors (TFs) that do not have known motifs. One direction of work is to expand our TF-target connections by leveraging expression-based network inference methods (De Smet & Marchal, 2010). Furthermore, much of our work relied on cell lines and primary tissues with available accessibility and histone modifications. Recent availability of single cell RNA-seq and accessibility datasets are greatly expanding our ability to define the diversity of cell types. Therefore, another direction of work would be to extend our approach to single cell omic datasets to both define long-range interactions, gene regulatory interactions, and identify cell type-specific gene subnetworks that are potential targets of regulatory variants. On the methodology side, richer representation learning approaches could be used to derive graph features for our graph clustering framework. Here, we relied largely on GWAS studies to define our set of variants. Another direction of work would be to consider a broader range of sequence variants, for example from whole genome sequencing of clinical phenotypes. As more genomes, transcriptomes and epigenomes of individuals and populations become available, approaches such as ours should be helpful to improve our ability to interpret non-coding variants and their impact on individual genes as well as on entire pathways.

## Materials and Methods

### L-HiC-Reg

L-HiC-Reg is based on HiC-Reg, a Random Forests based regression approach that predicts contact counts using one-dimensional regulatory genomic data sets, e.g. histone modifications and architectural proteins, and accessibility, (S. Zhang et al., 2019). L-HiC-Reg is specifically geared for cross-cell line predictions and was trained to use a smaller number of datasets compared to HiC-Reg. Furthermore, L-HiC-Reg uses discrete features (described below) compared to HiC-Reg which uses continuous features. To develop L-HiC-Reg, we first segmented a chromosome into 1MB bin and trained a Random Forests regression model for each adjacent 1MB segments using high-resolution (5kb) Hi-C SQRTVC normalized data downloaded from Rao et al (Rao et al., 2014). For a given 1MB segment, the training set included all 5kb region pairs in which at least one of the 5kb regions was inside the 1MB segment. We then made predictions on the same 1MB region in a test cell line. We trained Random Forests regression models for each of the human autosomal chromosomes. We concatenated predictions from multiple 1MB models for the entire chromosome, averaging predictions for pairs spanning two different adjacent regions.

### Feature Extraction and Pair Representation

#### Datasets

We used the following datasets to generate features for L-HiC-Reg: DNase1-seq, H3K27ac, H3K27me3, H3K36me3, H3K4me1, H3K4me3 and H3K9me3. We selected these signals because they were measured in the most cell lines and tissues in the Roadmap Epigenome Mapping consortium database. We obtained these datasets from the ENCODE project for the five cell lines with high resolution Hi-C data in Rao et al. K562, Gm12878, Hmec, Huvec and Nhek that we used for training and testing. Our test cell lines and their corresponding datasets came from the Roadmap database and included 55 cell lines that had measured DNase I profiles. However, only 22 cell lines had all these signals measured, therefore, we imputed signals in the absence of real signals as described below. In addition to the 7 datasets above, we used measured DNase I-seq to derive accessible binding motifs for CTCF, RAD21 and TBP, using PIQ (S. Roy et al., 2015; Sherwood et al., 2014) since ChIP-seq data is not available for these proteins in the Roadmap database. We obtained the raw fastq files from ENCODE (Dunham et al., 2012) and the Roadmap project (Roadmap Epigenomics Consortium et al., 2015) and aligned reads to the human hg19 assembly using bowtie2 (Langmead & Salzberg, 2012). Reads aligned to a locus were obtained using SAMtools (H. Li et al., 2009) and input to BEDTools (Quinlan & Hall, 2010) to obtain a base pair level read count which was then aggregated in a 5kb region. We then normalized the aggregated signal by sequencing depth and collapsed replicates by the median.

### Imputation of epigenomic signals

We generated imputations using a modified version of Avocado (Schreiber et al, 2020). Avocado is a deep tensor factorization method that organizes a compendium of functional data sets into a three-dimensional tensor with the axes being cell types, assays, and genomic positions. The learned latent representations for each axis, and the neural network predictor, are trained jointly using a mean-squared-error objective function on the available data sets. The trained model is then used to make predictions for all experiments, regardless of if they have corresponding experimental data. Avocado’s original formulation made predictions at 25bp resolution and, being multi-scale, had genomic latent representations at 25bp, 250bp, and 5kbp resolution. Because the data in this work was at 5kbp resolution, all latent representations were changed to be at 5kbp resolution. Other than that, Avocado was trained and used in the standard procedure as outlined in Schreiber et al, 2020.

### Feature extraction

L-HiC-Reg makes predictions for each pair of 5kb regions. As described previously (S. Zhang et al., 2019), we first generated features for each genomic 5kbp bin and represented each pair of regions using the features of the two regions and the window region between them. A ChIP-seq signal was represented as the average read count aggregated into a 5kb bin. To account for overall differences in signals across cell lines, we discretized each 5kb ChIP-seq signals using k-means clustering with k=20. For TBP, RAD21 and CTCF accessible motifs we used the number of motif instances with a PIQ purity score greater than 0.50 for each 5kb bin as features.

Each 5kbp region was represented as a 10-dimensional feature vector, each dimension corresponding to one of the 10 genome-wide datasets (6 histone ChIP-seq, DNase I-seq and the 3 DNase I-seq filtered motif instance counts). The feature value for the window region for a ChIP-seq or DNase 1-seq feature is the mean value of that signal after discretizing. For the motif count features, the window feature value is the number of motifs divided by the number of 5kb distance bins in the window. Each region in a given pair has a 10 dimensional feature vector, taken together with the feature vectors of the window region between the two regions to obtain a feature vector of size 30 (Whalen et al., 2016).

### Generation of contact counts in 55 Roadmap cell lines

We used the 5 trained L-HiC-Reg models using the Rao et al cell lines to generate the count predictions in the 55 cell lines. Our final analysis was done on the predictions from the model trained on the Gm12878 cell line, which is the cell line with the highest sequencing depth.

### Binomial method to call significant interactions

In order to determine significant interactions in the 55 Roadmap cell lines, we adopted the binomial method used in Duan et al., 2010 (Duan et al., 2010). The probability of observing a given interacting pair of 5kb bins exactly *k* times is computed via the binomial distribution:

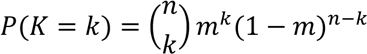

Where *m* is the probability of observing any interaction and is assumed to be uniform and *n* is the total number of observed interactions. Thus, the probability that each pair is observed at least *k* times is:

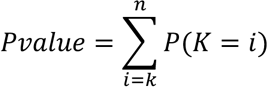

To control for the distance effect, we compute the p-values for every 5kb genomic distance bin up to 1MB. We also adopt the same multiple testing correction procedure as Duan et al., 2010, defining the q-value as the minimal false discovery rate threshold at which a given score is deemed significant.

## Evaluation Metrics

### Distance stratified Pearson’s correlation

To assess the quality of count prediction of L-HiC-Reg, we used Pearson’s correlation of predicted contact counts and true contact counts stratified by genomic distance as described previously (S. Zhang et al., 2019). This was done by grouping pairs of regions based on their genomic distance and calculating the Pearson’s correlation of predicted and true contact counts in each 5kb distance bin up to 1 MB. To easily compare the performance between L-HiC-Reg, HiC-Reg, transfer count and predictions generated from real and imputed data, we summarized the distance-stratified Pearson’s correlation curve into the Area Under the Curve (AUC) as described in Zhang et al (S. Zhang et al., 2019). Higher AUC indicates better performance.

### Identification of TADs and TAD similarity

To identify topologically associated domains (TADs), we applied the Directionality Index (DI) method described in (Dixon et al., 2012) using a genomic window of 2Mb on both real and predicted contact counts at a resolution of 5kb. We compare the similarity of TADs identified from the predicted counts (both HiC-Reg and L-HiC-Reg) and true counts for each chromosome and combination of train and test cell line as described previously in Zhang et al (S. Zhang et al., 2019). Briefly, we matched a TAD found in the true count data to a TAD found in the predicted counts based on the highest Jaccard coefficient. The Jaccard coefficients for each match were averaged across all TADs from the true counts. We repeated the process for each TAD from the predicted counts to a TAD in the true count and averaged the Jaccard coefficient across TADs from the predicted counts. The overall similarity between TADs was then the average of these two averages.

### Comparison of L-HiC-Reg predictions to computational and experimental approaches

We compared significant interactions from our compendium to two experimental datasets (1) cell line specific Capture Hi-C data (Jung et al., 2019) and (2) 10 ChIA-PET data sets (Heidari et al., 2014; G. Li et al., 2012). The Capture Hi-C experiments were performed on 27 cell lines, of which 9 overlapped with the cell lines for which we generated predictions. The ChIA-PET datasets included PolII in HeLa, K562 and Gm12878 cell lines, CTCF and TAD in the K562 and Gm12878 cell lines and multiple chromatin marks in K562 and Gm12878 cell lines (Heidari et al., 2014). We considered the ChIA-PET data to be non-cell line specific and tested all of the ChIA-PET datasets against all of the 55 cell lines.

Our metric for evaluating these genome-wide maps is fold enrichment, which compares the fraction of significant interactions identified from L-HiC-Reg that overlapped with experimentally detected measurements, against to the fraction of interactions expected by random chance. We mapped experimental (Capture Hi-C or ChIA-PET) interactions onto the pairs of regions used in L-HiC-Reg by requiring one region of an interaction from the experimental dataset to overlap with one region, and the other experimental region to map to the second region. Fold enrichment is defined as 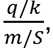, where *q* is the number of significant interactions from L-HiC-Reg that overlap an (/) interaction in the experimental data set, *k* is the total number of L-HiC-Reg significant interactions, *m* is the total number of interactions in the experimental dataset that can be mapped to any of the 5kbp pairs used for L-HiC-Reg, and *S* is the total number of possible pairs in the universe. A fold enrichment greater than 1 is considered enriched.

We also compared our results to two computational approaches (1) GeneHancer (Fishilevich et al., 2017), a non-cell line specific network that scores enhancer-promoter interactions based on various public datasets such as chromatin accessibility, eQTL and functional annotations and (2) JEME (Q. Cao et al., 2017), an approach that integrates chromatin accessibility, histone modifications and expression across a large number of biological samples to predict enhancer-promoter interactions within a binary classification setting. JEME first constructs a LASSO regression model around each promoter to identify candidate enhancers using expression and histone marks and then uses sample-specific errors and features to train a Random Forests model to predict enhancer-promoter interactions. We compared these methods to L-HiC-Reg using the fold enrichment metrics described above.

### Mapping significant interactions to genes and expression analysis

From our predictions in the 55 cell lines, we obtained significant interactions using the Binomial method (discussed above). We again mapped either end of significant interactions to genes using the Gencode v10 (2012) TSS annotation. A 5kb region in a pair can be associated with a gene if it overlapped within +/-2500bp of the TSS. Any gene that was associated to at least one significant interaction was considered “interacting” while any gene not mapped to any significant interaction was considered “non-interacting.” We downloaded RNA-Seq data from the Roadmap database which contained the RPKM values for 57 epigenomes. Of the 55 cell lines, 14 cell lines also had expression. We compare the log(RPKM) between interacting and non-interacting genes in these 14 cell lines using a T-test. We also performed this analysis for GeneHancer and JEME interactions.

### Mapping regulatory variants to genes with long-range interactions

We downloaded GWAS SNPs from the NHGRI-EBI GWAS catalog v1.0.2. We determined which GWAS SNPs were in non-coding regions based on whether they were annotated as “intergenic” or not. For all of these GWAS SNPs, we determined the SNPs that were in LD with the GWAS SNP with the tool LDproxy using an R^2 cutoff of 0.8. We identified significant interactions where the 5kb bin contains a coding gene promoter and the other bin contains a non-coding SNP across all of the 55 Roadmap cell lines.

### Mapping significant interactions to eQTLs

We used curl to query all GWAS and LD variants for 15 phenotypes in the GTEx v8 database (Lonsdale et al., 2013). This results in the gene symbols of all genes in eQTL with the variant. We evaluated the overlap between interactions from L-HiC-Reg, GeneHancer, and JEME with SNP-gene pairs in eQTL using (1) precision and (2) recall. We also compared to a baseline method, nearest neighbor (NN), which is the closest gene neighbor to a SNP by genomic distance. A SNP-gene pair were considered overlapping with the eQTL association if the SNP had a predicted interaction with the same gene. Predicted SNP-Gene pairs for L-HiC-Reg, JEME and GeneHancer were determined using the methods described in “Node Scoring” section of these methods. Precision is calculated as the total number of predicted interactions in eQTL over the total number of predicted interactions. Recall is calculated as the total number of predicted interactions in eQTL over the total number of eQTLs. These ratios are averaged over the phenotypes and cell lines.

### Examining the downstream effects of non-coding variants on cell line-specific gene networks

We developed a graph-theoretic framework to define gene subnetworks targeted by sets of SNPs. We first assembled cell line specific gene networks (**Network generation**). Nodes on each network were scored based on their long-range interactions to regulatory SNPs (**Node scoring**). The network nodes and edges were reweighted using graph diffusion (**Node and Edge Diffusion**) to define the combined effect of variants on genes based on the network structure. Finally, we applied Multi-task Graph Clustering to define groups of genes on the network based on their reweighted connectivity across all 55 cell lines.

### Network generation

We downloaded a protein-protein interaction network (PPI) from STRING version 9.1 (Mering et al., 2003). To obtain high-confidence edges, we considered only edges with a STRING confidence score > 0.95. To obtain cell line specific proximal networks, we downloaded motifs for 537 transcription factors from JASPAR (Khan et al., 2018). We applied PIQ to find accessible motifs using the Roadmap DNase I data in all 55 cell lines. We selected accessible motifs with a PIQ score greater than 0.9. We generated a TF-target regulatory network by mapping the accessible motifs to gene promoters using the Gencode v10 TSS annotations. Finally, we incorporated cell line specific, distal TF-target pairs via our long-range regulatory interactions. For a given cell line, we took the significant interactions with one region overlapping a gene TSS as defined by the Gencode v10 TSS annotations. We searched for a motif in the other region of the interacting pair using the same criteria as the proximal network. For both the proximal and distal networks, we took the top 5% of edges defined by the motif instance purity score. Finally, we merged the PPI network, and the proximal and distal TF-gene network to generate a cell-line specific network for each of the 55 Roadmap cell lines.

### Node scoring

For a given phenotype, we scored genes in a particular cell line based on whether they were connected to a non-coding GWAS SNP or an LD SNP via long-range interactions for that cell line. For each significant interaction, we matched one of the 5kb regions to a gene using the TSS (as described in “Mapping significant interactions to genes and expression analysis.”) and the other region to a SNP if its genomic coordinate was within the 5kb bin. The score of a gene was defined as the average -log(q-value) of all significant SNP-gene interactions involving that gene in a given cell line. Such genes were called “direct hits”. If a gene did not participate in a long-range interactions with SNPs, its score was set to 0.

### Node and Edge diffusion

The genes directly interacting with a SNP (direct hits) described above may not fully capture all of the downstream effects of non-coding variation of a given phenotype, which might be broader and facilitated by an underlying molecular network. Therefore, we applied a two-step graph diffusion approach. First, we applied node diffusion using the direct hit genes as input nodes. Node diffusion measures the influence of the input genes on all other genes in the network based on their global network connectivity. We used the regularized Laplacian kernel to estimate global network connectivity (Smola & Kondor, 2003). This kernel is defined as *K*_*L*_ = (1 + *λL*)^−1^ where, *L* is the symmetric normalized Laplacian and is defined as *L* = *I* − *D*^1/2^ *AD*^1/2^, where *A* is the adjacency matrix of the gene interaction network *G*, and *D* is a diagonal matrix giving the degree of each node. The diffusion score for a gene *g*_*i*_ is computed by *V* = *K*_*L*_ · *Q*, where *Q* is a 1-dimensional vector of input node scores and *V* is a vector of diffusion scores, (*v*_1_, *v*_2_, … *v*) where *n* is the total number of nodes in the graph. We tested different values of λ, {0.1, 0.5, 1, 5, 10}, which is a hyper-parameter and specifies the kernel width, using leave-one-out cross-validation. We tested these values for lambda for all 55 cell lines and several phenotypes (inflammatory skin disease, inflammatory bowel disease, resting heart rate and pediatric autoimmune disease) using a leave one out cross-validation approach. Treating the direct gene hits as labeled data, we scored each value of λ using the area under the precision-recall curve (AUPR) computed from leave one out cross-validation. For each fold, we left one of the direct hits at a time, carried out diffusion and used the diffusion score when it was not included in the input set to predict its label. At a diffusion score *v*_*i*_, precision is computed as the fraction of all genes with score ≥ *V*_*i*_, that are in the query set, and recall is computed as the fraction of all the genes in the query set that have score ≥ *V*_*i*_. In general, λ ≤ 1 performs the best across cell lines and tissues, with minimal difference in AUPR between λ = 0.1,0.5, 1. λ = 10 rarely performs the best. Therefore, we decided to use λ = 1 across cell lines and phenotypes. Following node diffusion, we reweighted the edges using edge diffusion to obtain cell-type specific weighted graphs. We used an insulated heat diffusion kernel (Vandin et al., 2011)(Leiserson et al., 2015) to estimate the influence of a node on its neighboring nodes based on their global connectivity. The insulated heat kernel is defined as *K*_-_ = β(*I* − (1 − β)*W*)^−1^. *W*is a transition matrix defined as *W* = *AD*^−1^, where *A* is the adjacency matrix of the gene interaction network, and *D* is a diagonal matrix of node degrees. The parameter *β* specifies the retention rate of the kernel and was set to a default value of 1 in this study. We define a diagonal matrix *D*., where an element of the matrix *d*_(*i,i*)_ is the diffused nodes score *V*_*i*_ from the node diffusion step. We computed the final weighted adjacency matrix as *H* = *K*_*H*_ · *D*_*V*_. This matrix is converted into a diffusion-state distance (DSD) matrix (M. Cao et al., 2014), which was shown to improve the ability to detect gene modules on graphs (Choobdar et al., 2019). Briefly, we defined a *n*-dimensional vector of the diffusion state of a node *g*_*i*_ as *u*_*i*_ = (*h*_(*i,1*)_, *h*_(*i,2*)_, …, *h*_(*i,n*)_ from each row of *H*. The DSD matrix *P* was defined by the elements *p*_(*i,j*)_ = ||*u*_*i*_ − *u*_*j*_||_1_. After obtaining the DSD matrix, we converted it into a similarity matrix *S* via a Gaussian kernel defined as *S* = *exp*(−*P*^2^/2σ^2^_*P*_), where σ_*P*_ is a standard deviation of all elements of the matrix *P*. The final result is a cell type specific reweighted network for each of the 55 cell lines where the weights correspond to the similarity values of the diffusion profiles of the nodes.

### Multi-task graph clustering

To identify subnetworks in each of the cell lines, we applied a multi-task graph-based clustering algorithm, Muscari (Shin et al., 2021). This algorithm finds the network clusters in multiple cell types by simultaneously applying spectral clustering to each cell type-specific network while incorporating the relatedness of the cell types. A key property of this multi-task learning framework is that there is a mapping of clusters from one cell line to another. Therefore, cluster *i* in one cell line corresponds to cluster *i* in another cell line. For spectral clustering of each cell line-specific network, we used eigenvector matrices of the regularized graph Laplacian, *L*_*τ*_ = *I*− *D*^1/2^_*τ*_ *AD*^1/2^_*τ*_, where *A* is the adjacency matrix and *D*_4_ is a regularized diagonal matrix defined as *D*_*τ*_ = *D* + μ_*D*_, where μ_*D*_ is a mean of *D* (Y. Zhang & Rohe, 2018).

Muscari takes as input the number of clusters in each cell line, input graphs and the relationship between the cell lines. We ran Muscari with k = {6,7,8,9}, and selected k=7 based on modularity. We used hierarchical clustering of H3K4me3 signal in promoters as the measure of relatedness for the 55 cell lines and tissues. Briefly, we created an *N* x *M* matrix of 5kb-aggregated H3K4me3 ChIP-seq signals, where *N* is the number of cell lines and *M* is the total number of promoter-associated 5kb bins throughout the entire genome. We performed hierarchical clustering with Euclidean distance using unweighted average distance (UPGMA) for computing the distance between clusters. As a basis for comparison, we also clustered the Roadmap cell lines by their shared 3D genome conformation. For every possible pair of Roadmap cell lines, we considered the significant interactions from each cell line as a network and compared the two networks using the F-score. The F-score matrix was converted into a distance matrix based on Euclidean distance between pairs of cell lines of their overall F-scores. This was used as input to hierarchical clustering. Hierarchies learned from both metrics had similar structure as assessed using the Fowlkes-Mallows index (**Supp Figure 3**).

### GO enrichment on Muscari clusters

We performed GO enrichment on each cluster separately using a hypergeometric test with FDR correction (FDR < 0.05) and all genes in the phenotype as the background. As the number of gene lists (cluster in a cell line) we tested for enrichment was substantial (55 x 7), we applied a matrix factorization approach to identify groups of terms for groups of gene lists to enable easier interpretation. We first merged the GO terms into a matrix of -log(P) values, *X*, that was *N x M*, where *N* is the total number of terms across the 7 clusters in any cell line and *M* is all cell lines in all clusters (55 x 7). On this matrix, we performed Non-negative Matrix Factorization (NMF) with k=7 and used the resulting row and column factors to cluster the terms and lists. For each NMF cluster, we took the top 10 terms and the top 5 cell lines based on their values in the lower dimensional factors. We present this result in **Supp Figure 5**.

### Identification of dynamic gene sets

After applying Muscari to all 55 cell lines, we obtain cluster assignments for all genes in the network. A dynamic gene set is a gene set with a similar cluster membership in one part of the tree of cell lines, but different membership, but similar across genes, in another part of the tree. We identified these gene sets based on the *de novo* clustering approach described in (Shin et al., 2021) on Muscari outputs. Briefly, hierarchical clustering was performed on the cluster assignment profiles of genes followed by optimal leaf ordering and then gene sets were obtained with a cut height of 0.1. We removed gene sets with less than 5 genes.

We interpreted these gene sets based on overlap of genes with SNPs, tendency of SNP-gene interactions to vary according to the change in cluster assignment, and Gene Ontology enrichment.

### Data availability

Predicted significant interactions, SNP-Gene interactions, node and edge scores, networks, transitioning gene sets and all code associated with this project can be found at: https://github.com/Roy-lab/Roadmap_RegulatoryVariation.

## Acknowledgements

We would like to thank Erika Da-Inn Lee for assistance with visualizations for this project. We would also like to thank the Center for High-Throughput Computing (CHTC) at UW-Madison for providing resources to complete this project. This work was supported by the Genomics Sciences Training Program at UW-Madison (NHGRI 5T32HG002760) for BB, NHGRI R01 grant R01- HG010045-01 for SR and BB, the Center for Predictive Computational Phenotyping (NIH BD2K U54 AI117924) for BB and SR, and James McDonell Foundation Grant 3194-133-349500-4- AAB5159 for SR and JS.

