## Supplemental Figures for "Leveraging epigenomes and three-dimensional genome organization for interpreting regulatory variation"

### Supplementary figures

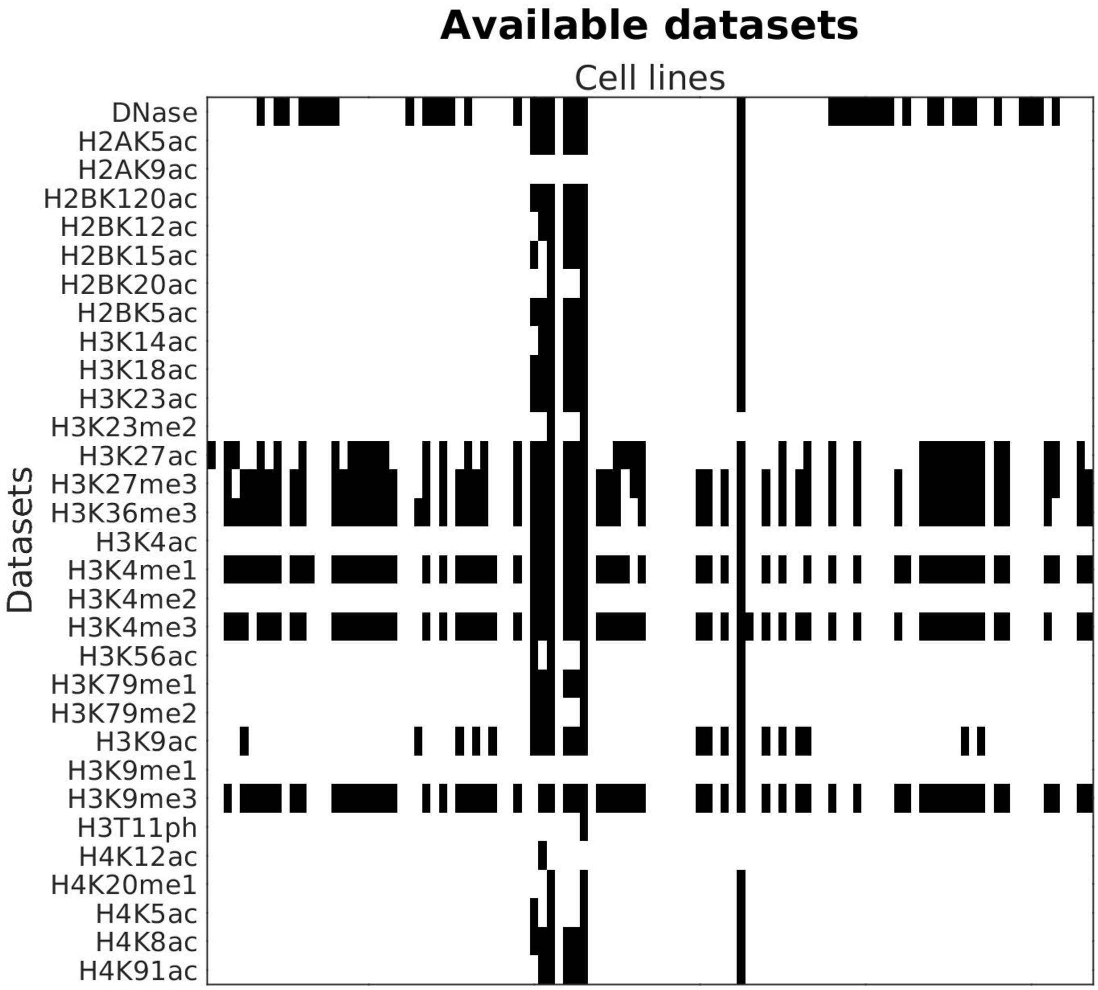


**Supplemental Figure 1.** Datasets available from the Roadmap epigenomics database across cell lines (columns). Black – dataset exists in Roadmap, White – dataset absent from Roadmap.

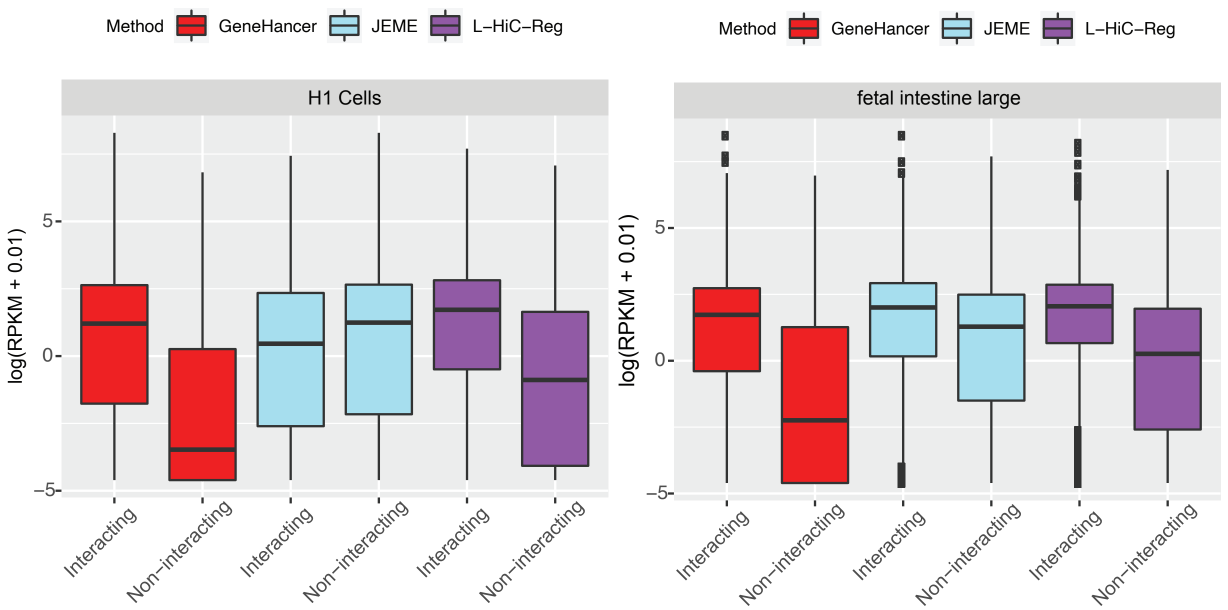


**Supplemental Figure 2.** Expression of interacting versus non-interacting genes for additional cell lines with expression data and predictions in both JEME and L-HiC-Reg. Related to Main Figure 3.

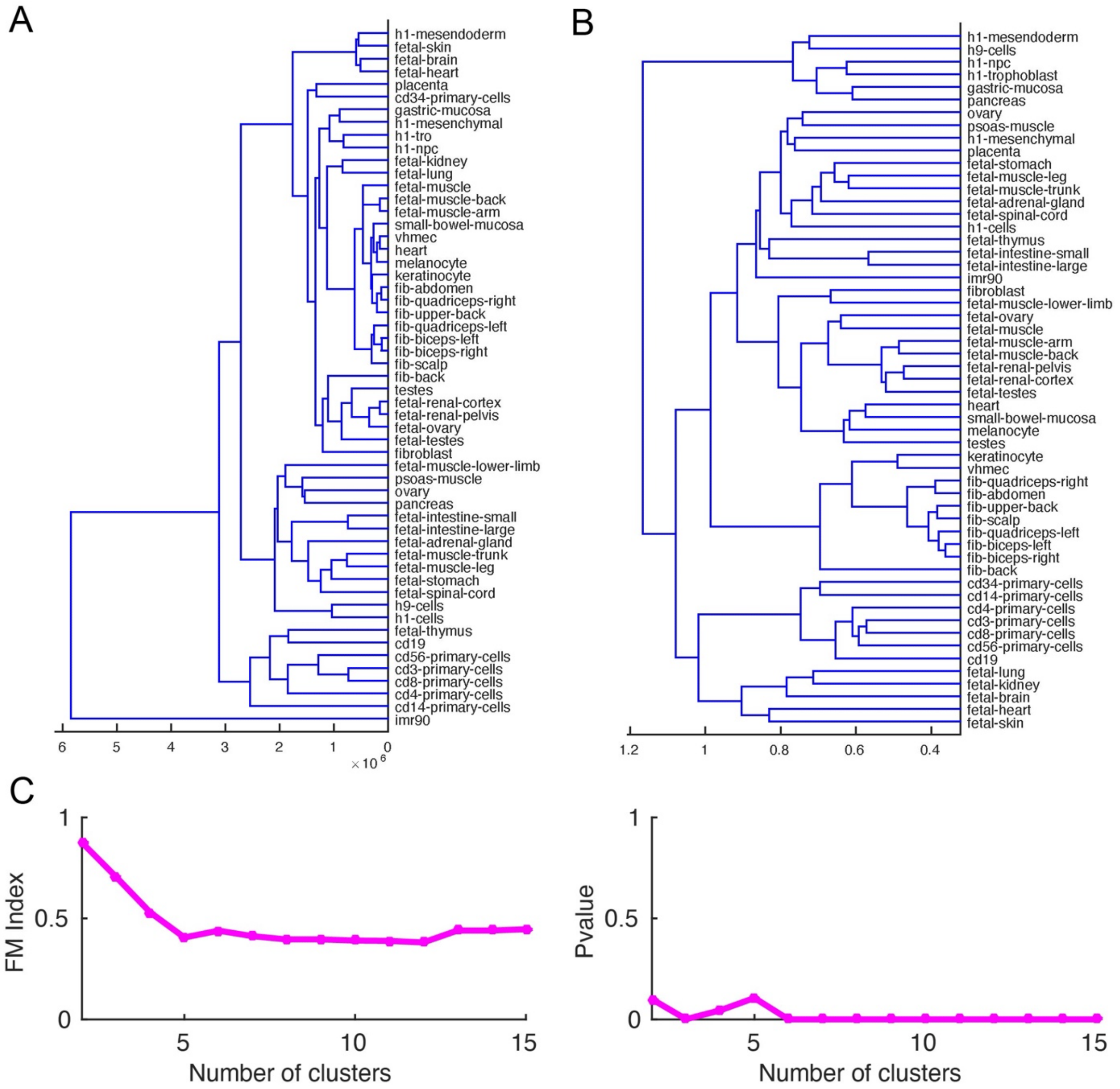


**Supplemental Figure 3. A.** Hierarchical clustering tree showing the similarity of H3K4me3 ChIP-Seq signal in the TSS of genes. This tree was used as input for the Multi-task Graph Clustering algorithm. **B.** Hierarchical clustering based on the F-score of shared interactions between cell lines. **C.** Fowlkes-Mallows index (left) comparing the similarity of two hierarchical clusterings at different levels of clustering with the associated P-value (right) based on random permutation.

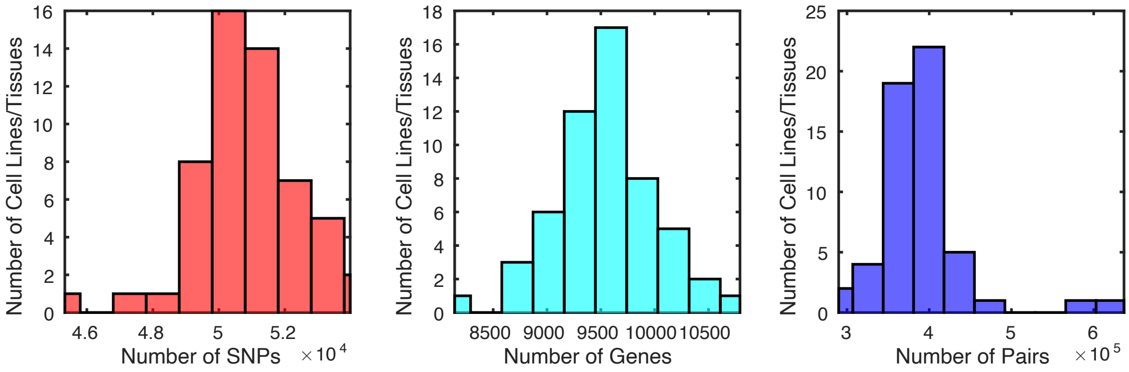


**Supplemental Figure 4.** Distribution of the number of SNPs in LD with the GWAS non-coding SNPs from the NHGRI-EBI GWAS Catalog (left), the number of genes mapped to these SNPs (middle) and the number of pairs connect SNPs to genes across multiple cell lines (left).

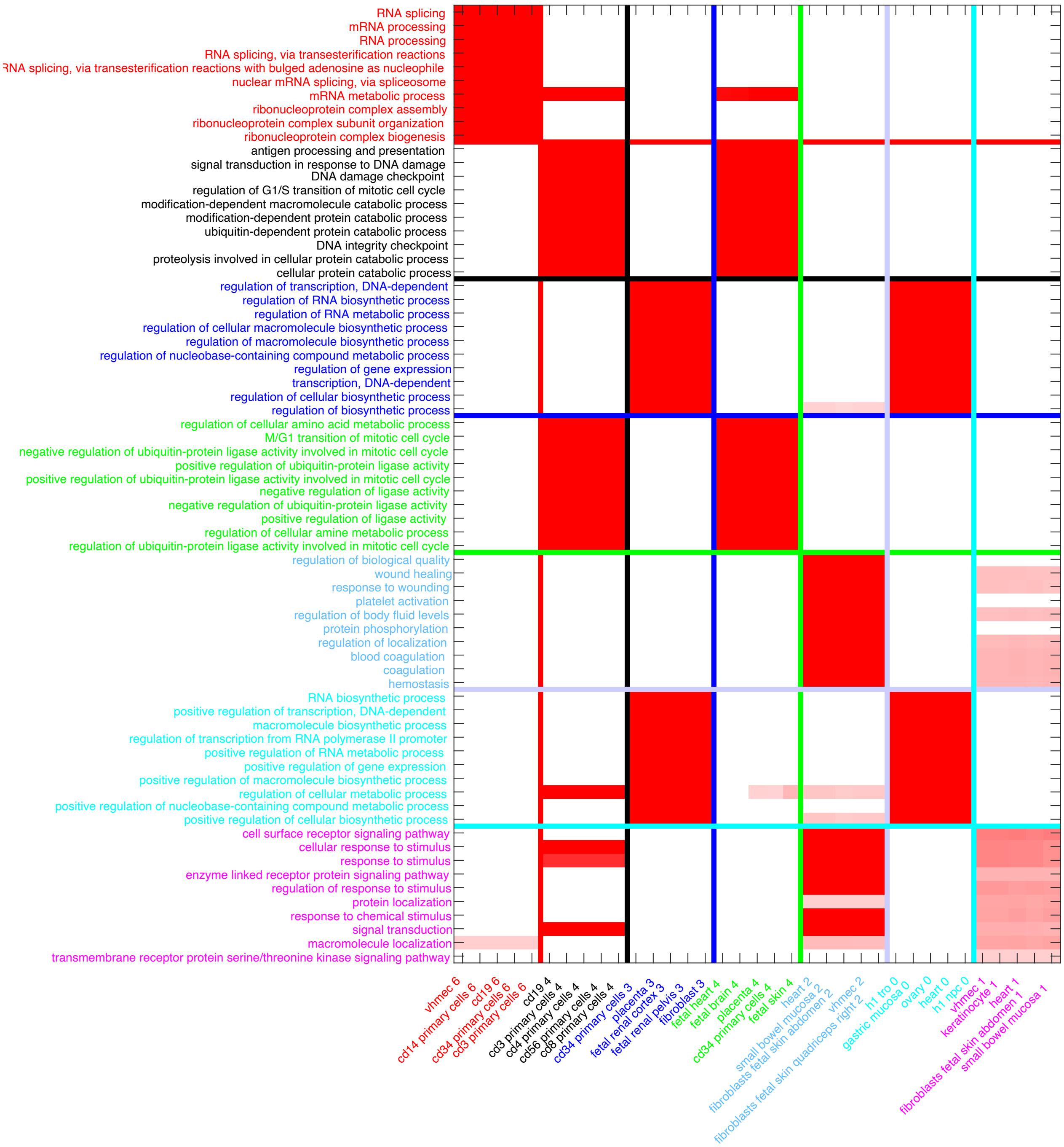


**Supplemental Figure 5.** Top Gene Ontology terms and cell lines across the 7 multi-task graph clustering clusters. Numbers next to the cell line name represents the graph cluster. To enable extraction of meaningful annotation from our GO analysis, we applied Non-negative Matrix Factorization with seven low-dimensional factors on the -log(q-value) scores of all the GO terms across all cell lines and gene clusters. For each NMF factor, the top 10 GO terms and the top 5 cell lines were selected based on their values in the corresponding lower-dimensional factors.

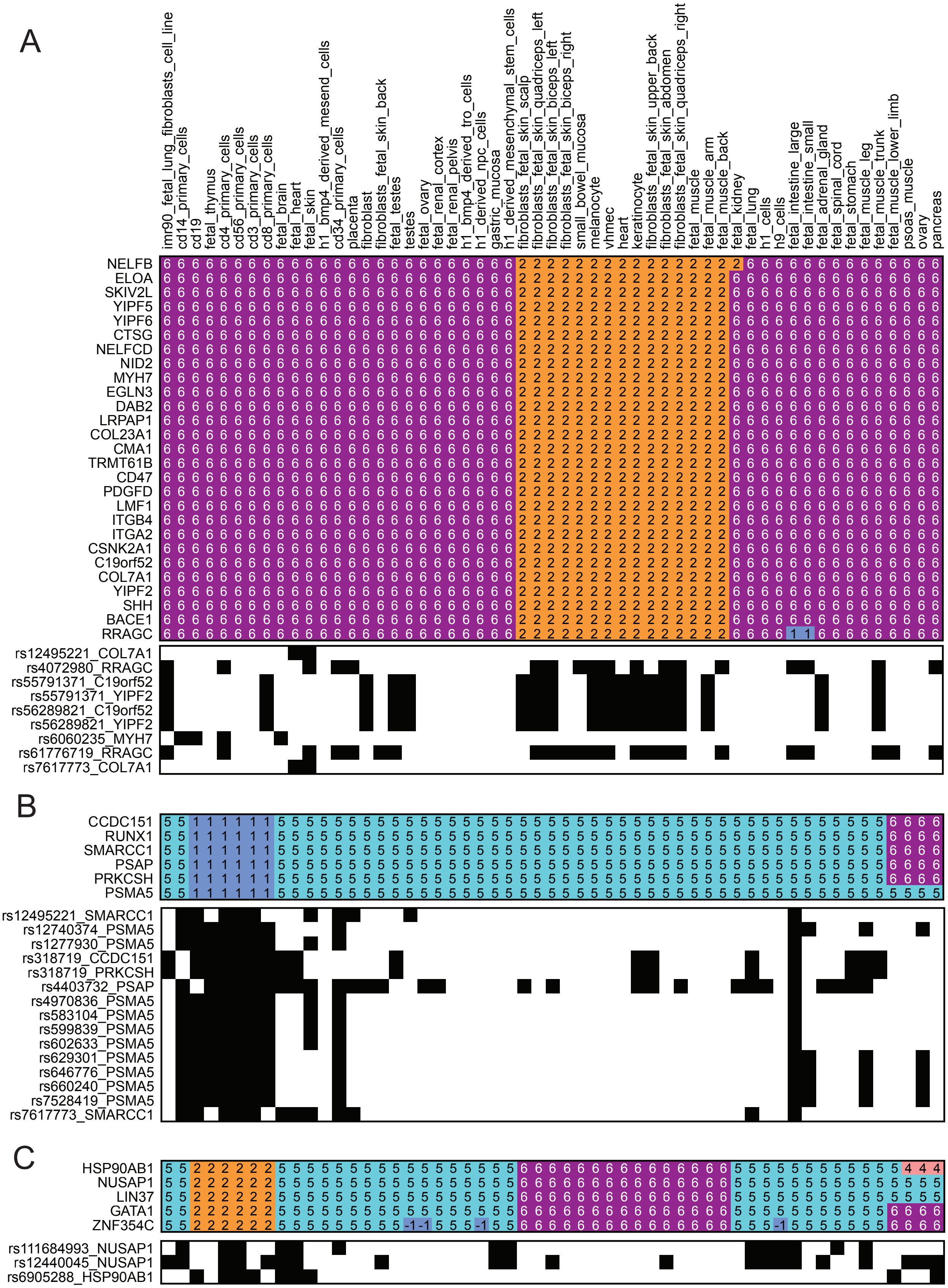
A
imr90_fetal_lung_fibroblasts_cell_line
cd14_primary_cells
cd19
fetal_thymus
cd4_primary_cells
cd56_primary_cells
cd3_primary_cells
cd8_primary_cells
fetal_brain
fetal_heart
fetal_skin
h1_bmp4_derived_mesend_cells
cd34_primary_cells
placenta
fibroblast
fibroblasts_fetal_skin_back
fetal_testes
testes
fetal_ovary
fetal_renal_cortex
fetal_renal_pelvis
h1_bmp4_derived_tro_cells
h1_derived_npc_cells
gastric_mucosa
h1_derived_mesenchymal_stem_cells
fibroblasts_fetal_skin_scalp
fibroblasts_fetal_skin_quadriceps_left
fibroblasts_fetal_skin_biceps_left
fibroblasts_fetal_skin_biceps_right
small_bowel_mucosa
melanocyte
vhmec
heart
keratinocyte
fibroblasts_fetal_skin_upper_back
fibroblasts_fetal_skin_abdomen
fibroblasts_fetal_skin_quadriceps_right
fetal_muscle
fetal_muscle_arm
fetal_muscle_back
fetal_kidney
fetal_lung
h1_cells
h9_cells
fetal_intestine_large
fetal_intestine_small
fetal_adrenal_gland
fetal_spinal_cord
fetal_stomach
fetal_muscle_leg
fetal_muscle_trunk
fetal_muscle_lower_limb
psoas_muscle
ovary
pancreas
NELFB
ELOA
SKIV2L
YIPF5
YIPF6
CTSG
NELFCD
NID2
MYH7
EGLN3
DAB2
LRPAP1
COL23A1
CMA1
TRMT61B
CD47
PDGFD
LMF1
ITGB4
ITGA2
CSNK2A1
C19orf52
COL7A1
YIPF2
SHH
BACE1
RRAGC
rs12495221_COL7A1
rs4072980_RRAGC
rs55791371_C19orf52
rs55791371_YIPF2
rs56289821_C19orf52
rs56289821_YIPF2
rs6060235_MYH7
rs61776719_RRAGC
rs7617773_COL7A1
B
CCDC151
RUNX1
SMARCC1
PSAP
PRKCSH
PSMA5
rs12495221_SMARCC1
rs12740374_PSMA5
rs1277930_PSMA5
rs318719_CCDC151
rs318719_PRKCSH
rs4403732_PSAP
rs4970836_PSMA5
rs583104_PSMA5
rs599839_PSMA5
rs602633_PSMA5
rs629301_PSMA5
rs646776_PSMA5
rs660240_PSMA5
rs7528419_PSMA5
rs7617773_SMARCC1
C
HSP90AB1
NUSAP1
LIN37
GATA1
ZNF354C
rs111684993_NUSAP1
rs12440045_NUSAP1
rs6905288_HSP90AB1

**Supplemental Figure 6.** Gene sets with changes in graph cluster assignment across cell lines that exhibit concordant presence-absence of SNP-gene interactions for Coronary Artery Disease (CAD). Shown are three gene sets, #37 (**A.**), #46 (**B**.) and #53 (**C.**). Color heatmaps represent the cluster assignments for all 55 cell lines and the black and white heatmaps represents the presence (black) or absence (white) of the SNP-gene interaction with specific genes in the gene set across the cell lines.
